# Cadherins modulate the self-organizing potential of gastruloids

**DOI:** 10.1101/2023.11.22.568291

**Authors:** Alexandre Mayran, Dominique Kolly, Lucille Lopez-Delisle, Yuliia Romaniuk, Maxine Leonardi, Anne-Catherine Cossy, Theo Lacroix, Ana Rita Amândio, Pierre Osteil, Denis Duboule

## Abstract

Gastruloids have recently emerged as an efficient four-dimensional model for studying some aspects of post-implantation embryonic patterning. They undergo gastrulation-like processes leading to the self-organization into highly reproducible biological objects. Here, we sought to uncover the molecular and cellular mechanism underlying this remarkable property. We report that self-organization competence is associated with a cell-specific coordination of a Cadherin switch. We find that N-Cadherin hinders gastruloids morphogenetic competence, for its inactivation leads to the formation of trunk-like structures in absence of extra-cellular matrix analogues. In contrast, *E-Cadherin* repression by *Snai1* is critical for self-organization: *Snai1* establishes a cell-specific repressive pace by triggering the repression of a pluripotency-associated transcription program and its chromatin landscape, thus allowing a proper transition from E-to N-Cadherin to occur. Altogether, this work establishes a molecular mechanism that integrates the exit from pluripotency and the pace of cell differentiation, leading to the observed self-organizing potential of gastruloids.

## INTRODUCTION

Early vertebrate embryos can display a wide range of morphologies and organisations, depending on environmental conditions. Species-specific adaptations develop on top of a conserved vertebrate archetype. Indeed, at late gastrulation, the various embryonic body plans show remarkable consistencies^1^. This rather generic developmental stage is pivotal for the embryo and relies on a combination of coordinated cell movements, morphogenesis and cell fate specification to organize the three germ layers (ectoderm, mesoderm and endoderm) along the emerging embryonic axes^2^. This process relies on signalling and mechanical cues arising from the extra-embryonic tissues, which instruct where gastrulation initiates. At the primitive streak, a region defining the posterior epiblast will undergo epithelial to mesenchymal transition (EMT), in response to BMP, WNT and Nodal signalling emanating from extra-embryonic ectoderm, thus leading to cell ingression and the formation of the future mesodermal layer. In contrast, the anterior epiblast will respond to antagonist BMP, Nodal and WNT signalling from the anterior visceral endoderm preventing streak formation, which will maintain an epiblast identity and form the embryonic ectoderm. These interactions between the extra-embryonic and embryonic compartments are thought to be essential for proper patterning of the embryo^2–4^.

The recent development of various *in vitro* stem cell-derived models of embryos can provide insight into these complex mechanisms. To some extent, these paradigms can recapitulate the development either of blastocysts and advanced pre-implantation embryos by combining trophoblast stem cells (TSCs) with embryonic stem cells (ESCs)^5^, or of post-implantation embryos by also including extra-embryonic endoderm cells (XEN)^6,7^. Amongst other properties, these models represent a useful minimal system to investigate the communication between embryonic and extra-embryonic tissue. However, the aggregation of mouse ESCs alone can also be induced to generate distinct embryonic-like structures referred to as gastruloids^8–10^. They can organize polarities and elongate an anterior-posterior axis in the absence of any extra-embryonic tissue and in a highly reproducible manner^8–12^, which in many respects mimics tail bud elongation, thus revealing the previously under-appreciated ability of embryonic cells to form complex embryonic-like structures with minimal pre-existing positional cues^13^. This ability which we refer to as self-organisation can be triggered by various phenomenon; for example a recent study^14^ identified an early radial symmetry breaking determining a binary response to WNT. Indeed, since gastruloids are initiated as a sphere, cells from the outer portion could have different mechanical, signalling, nutrient, aerobic status than cell deeper in the sphere. Alternatively, this radial symmetry breaking may be a consequence of a pre-existing heterogeneity in cellular state which then self-organize into their outer-inner portions. Finally, self-organisation may also arise from a cell-sorting mechanism of a disordered initial structure. For example, differential cell adhesion properties between endodermal cells and the remaining of gastruloid lineages was recently proposed to participate in the spatial organisation of endoderm in gastruloids^15,16^. Altogether, an integrated view of the molecular, cellular and tissue scale of self-organisation is nevertheless still lacking.

While complex spatio-temporal patterns of gene expression are observed in gastruloids, they lack the associated morphologies such as for example segmented somites surrounding a neural tube. However, by providing extra-cellular matrix (ECM) components (Matrigel), they can form these missing structures^17,18^, illustrating the value of this model to investigate the minimal requirement for some embryonic processes to occur. Furthermore, gastruloids can also be generated either from human pluripotent stem cells^19,20^ or from dissociated zebrafish embryo explants^21^ with a similar absence of defined morphologies and the same ability to self-organize and elongate an axis, suggesting that the latter property relies upon a vertebrate ancestral program. While the symmetry-breaking and self-organizing processes are being intensely studied^14,15,22,23^, the nature of the underlying molecular and genetic mechanisms remains elusive. In this study, we aimed to dissect the chain of events allowing gastruloid self-organization to integrate the molecular, cellular and tissue level scale. We identified an integrated process taking gastruloids through this robust phase of self-organization, as well as its association with cell fate transitions. This is achieved by Snai1, which induces the cell-specific repression of a pluripotency program including E-Cadherin, setting the pace of the differentiation and triggering appropriate lineage segregation. In contrast, the inactivation of N-Cadherin is sufficient to increase the self-organizing potential and generate highly reproducible structures with evidence of somite patterning surrounding a neural tube, overcoming the requirement for ECM.

## RESULTS AND DISCUSSION

### Gastruloid self-organization is resilient to intrinsic and extrinsic variations

Gastruloids develop axially organised structures composed of cells derived from all three germ layers^10,16,18^. Following a pulse of exogeneous WNT activation (Chiron, CHIR99021), they break symmetry and elongate an antero-posterior axis^8–10^ without the influence of extra-embryonic tissue or from the implantation into the uterine wall. This self-organization capacity may involve distinct cell signalling, mechanical constraints between inner and outer cells, or a differential accessibility to Chiron, as well as heterogeneity in the starting cellular state. To limit this latter point, gastruloids were generated from ESCs grown in a medium containing GSK3i and MAPKi (referred to as 2i) where ESCs cultures is more homogeneous^11,14,16,24^ (Fig. S1A, B).

First, we sought to disrupt cell positional information during gastruloid development, to evaluate whether it is required for gastruloid self-organisation. We performed single-cell dissociation of gastruloids at different developmental stages and re-aggregated them. When dissociated either at 48h, or at 72h and re-aggregated, gastruloids developed as control specimens, yet with a smaller size caused by some cells being lost along the procedure (Fig. 1A, Supplementary movie 1, Supplementary note 1). This suggests that at these early stages, there are no critical positional information and that outer and inner cells can be reshuffled without impacting the final structure. In contrast, when the already asymmetric 96h gastruloids were dissociated and re-aggregated, proper elongation and patterning were not observed, suggesting that the self-organization capacity is lost between 72h and 96h.

**Fig. 1:**
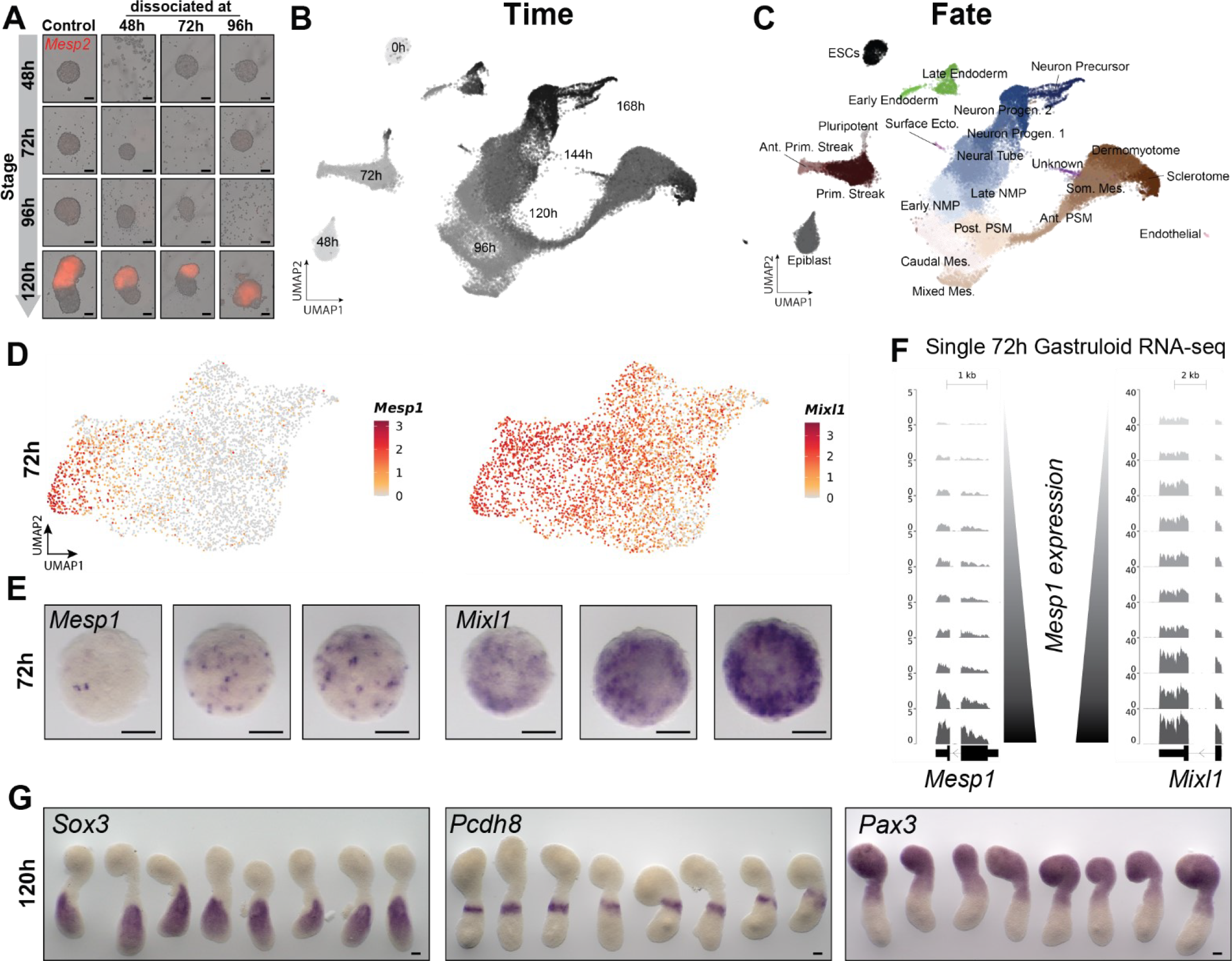
Gastruloid self-organization competence is resilient to intrinsic variation. (**A**) Representative image of gastruloids from a Mesp2^mCherry^ reporter line at the indicated stages. Rows identify the timing of gastruloid dissociation. **(B, C)** UMAP plot coloured by time of gastruloids development **(B)** or by clusters **(C)** of single cell RNA-seq of ∼52’000 gastruloids cells from the indicated time points. Two to five replicates were generated for each sample from 72h to 168h. ESCs, Embryonic Stem Cells; Prim. Streak, Primitive Streak; Ant., Anterior; Mes., Mesoderm; NMPs, Neuro-Mesodermal Progenitors; Progen., Progenitor; Surf. Ecto, Surface Ectoderm. **(D)** UMAP plot of 72h gastruloid cells showing the expression of *Mesp1* and *Mixl1* markers of the anterior primitive streak fate at 72h. **(E)** *In situ* hybridisation for anterior primitive streak markers *Mesp1* and *Mixl1*, at 72h. **(F)** Normalised coverage (read per million) of single gastruloids RNA-seq from GSE106227 showing *Mesp1* and *Mixl1* expression. Gastruloids are ranked by increasing level of *Mesp1* expression (FPKM). **(G)** *In situ* hybridisation for marker of neuronal (*Sox3*), anterior pre-somitic mesoderm (*Pcdh8*) and somitic mesoderm (*Pax3*) identities on 120h gastruloids. For the whole figure, scale bars are 100 μm.

We then assessed the link between self-organisation competence and cell identity. We documented the emergence and extent of cell diversification by conducting single cell RNA sequencing analysis of 52’000 gastruloids cells spanning from 48h to 168h, with a 24h resolution, also including the starting ESCs population as the 0h time-point (Fig. 1B, C). At 48h, all cells exhibited an epiblast identity and no transcriptional diversity was detected. (Fig. 1B, C, Fig. S1C, D). This suggests that the ability to self-organize does not rely on a pre-existing heterogeneity of cellular states. Then, at 72h, a global response to WNT signalling was visible with primitive streak (PS)-like cells expressing *Brachyury* and *Fst* (Fig. S1D, E). In addition, cellular diversification was initiated (Fig. 1D, Fig. S2A, B) and an anterior primitive streak (APS) program was expressed in 15% of gastruloid cells. At that stage, gastruloids were still fully able to self-organise even following dissociation (Fig. 1A) which means that cell fate diversification precedes the loss of self-organization competence. These APS expressing cells showed a salt and pepper distribution, with a variable number of positive cells from one gastruloid to the next (Fig. 1E, Fig. S2C). When comparing gene expression between single gastruloids^10^, APS-program genes correlated well with each other (Fig. 1F, Fig. S2D), as was also observed in the single cell transcriptome dataset (Fig. S2E). This supported a stochastic activation of this transcriptional program as a whole, rather than gene-specific stochastic transcriptional noise. Since the known regulators of this gene network, *Mixl1* and *Eomes*^25–27^, were the most extensively expressed (Fig. 1D, Fig S2B), they likely trigger the activation of this program.

Advanced gastruloids (from 96h), contain derivatives from all three germ layers, with mesodermal, neuronal, endodermal, endothelium and PGC/Pluripotent cells (Fig 1C, Fig S1D, E)^17,18^. Despite the variable nature of the earliest cell diversification event at 72h, the spatial organisation of elongated gastruloids at 120h was highly reproducible (Fig. 1G, Fig. S2G). NMPs marker genes (*Cyp26a1, Cdx2, Hoxb9*) were expressed at the posterior aspect of gastruloids, as expected^17,18^. Also, anterior progression to more differentiated status was observed for both neuronal and somitic lineages. Of note, few tested posterior markers showed inconsistent patterns such as the pre-somitic mesoderm marker *Tbx6*, the EMT regulator *Snai1* or the floor plate marker *Shh*. In these latter cases, however, expression was never scored in anterior gastruloids (Fig. S2H).

From this analysis, we concluded that gastruloids ability to self-organize is constrained in time and that the critical positional information required for the late gastruloid pattern is only established at ca. 96h. The gastruloids ability to form robust patterns of gene expression, despite an intrinsic early variability is thus consistent with a cell-sorting mechanism^15,23^.

### Cell specific coordination of a Cadherin switch

Amongst important mediators of cellular sorting and segregation are Cadherins^28^. Also, during gastrulation, part of the epiblast undergoes an EMT, which is characterised by a switch from E to N-Cadherin expression. Since gastruloids cells at 48h globally resemble epiblast (Fig. 1, Fig S1), we assessed whether a comparable E-to-N Cadherin switch was observed during gastruloid development (Fig. 2A). At both 48h and 72h, gastruloid cells expressed E-Cadherin (*Cdh1*), whereas N-Cadherin (*Cdh2*) was expressed in most cells from 96h to 168h, except in the PGC/pluripotent population. Of note, the initiation of *Cdh2* expression occurred at 72h within the APS population, in cells still expressing *Cdh1*.

**Fig. 2:**
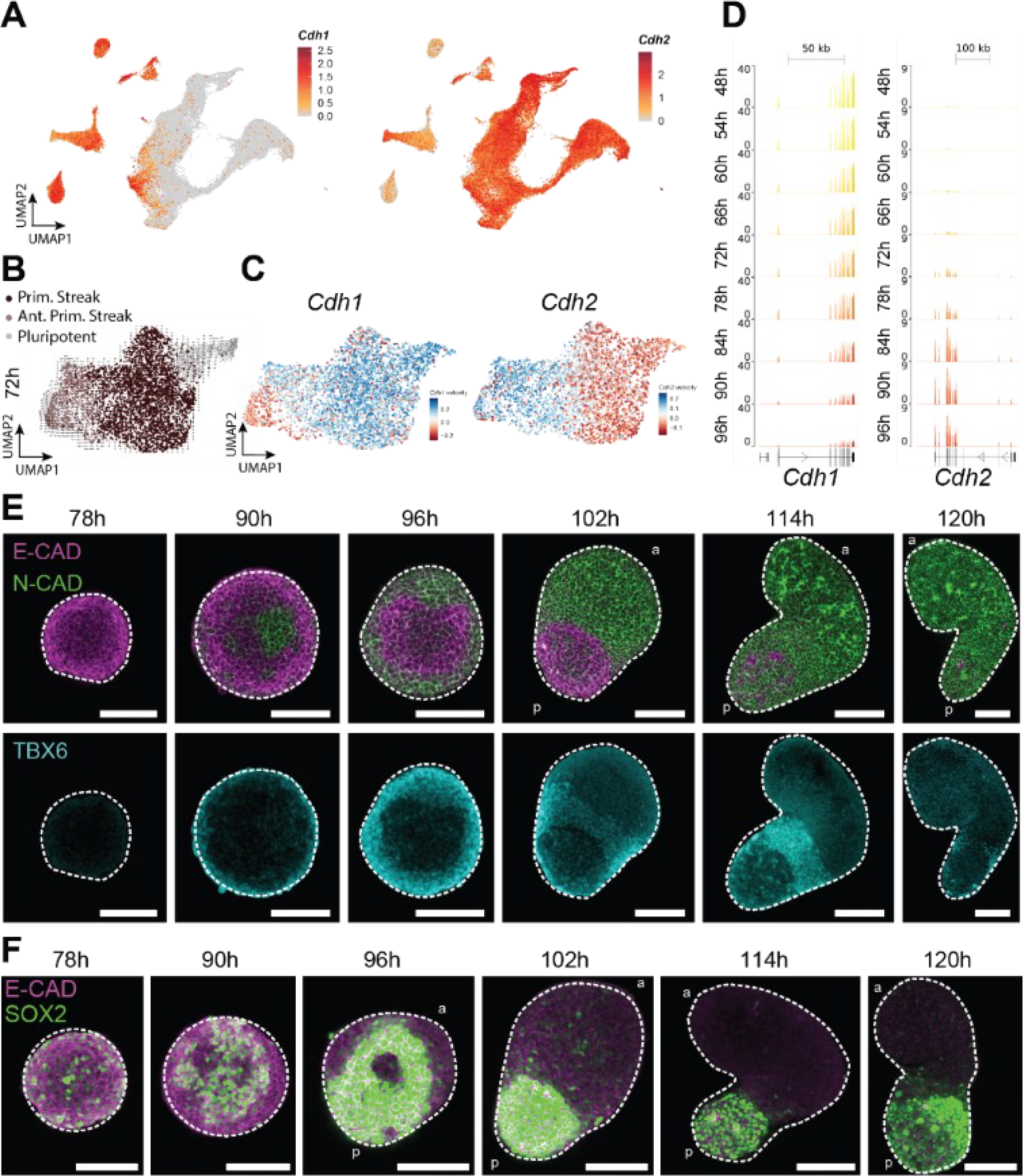
Cell-specific Coordination of a Cadherin switch is linked to self-organisation. **(A)** UMAP plot coloured by normalised expression of *Cdh1* (E-Cadherin) and *Cdh2* (N-Cadherin) across gastruloid stages. **(B)** UMAP plot of 72h gastruloids, coloured by cluster identity arrows represent the computed RNA velocity vectors^59^. **(C)** UMAP plot of 72h gastruloids, coloured by gene-specific RNA velocity on *Cdh1* and *Cdh2*. **(D)** Normalised coverage from bulk RNA-seq (read per million) at the *Cdh1* and *Cdh2* locus at the indicated time points (48h to 96h every 6 hours), average of three replicates. **(E-F)** Immuno-fluorescence for E-Cadherin, N-Cadherin (Top panels) and TBX6 (Bottom panels) (**E**), and E-Cadherin and SOX2(**F**)on gastruloids at the indicated time points. Scale bars are all 100 μm, at least 5 gastruloids were imaged on a confocal microscope Sp8 and the stack is 20 μm deep into the gastruloid for each image. For each condition where anterior posterior is identifiable, “a” designate the anterior side and “p” designate the posterior side. Gastruloids contours are represented by the dashed line.

We characterised the transcriptional dynamic of this transition using RNA velocity (Fig. 2B) and the activation of *Cdh2* occurred in a large proportion of 72h gastruloids cells. In contrast, a smaller percentage of APS cells had started to switch off *Cdh1* (Fig. 2C). These same cells also showed the activation of *Snai1*, a EMT regulator known to repress E-Cadherin expression (Fig. S3A). This suggests that N-Cadherin activation precedes E-Cadherin repression during gastruloid development.

To precisely establish the temporal sequence of the E-to N-Cadherin switch, we performed a comprehensive transcriptome time-series between 48h to 96h, with a 6h resolution (Fig. S3C). At 72h, gastruloid cells first switch on *Cdh2* expression, with a maximum already reached at 84h. Instead, *Cdh1* repression was only detected at 84h and expression decreased until 96h (Fig. 2D). We next assessed the spatio-temporal dynamics of E- and N-Cadherin proteins using immuno-fluorescence (Fig. 2E) and quantified their membrane level of expression (Fig. S4A). At 78h, virtually all cells contained E-Cadherin, while N-Cadherin was not detected, suggesting a delay between the transcripts and protein dynamics. By contrast, widespread expression of N-Cadherin was scored at 120h while E-Cadherin was detected scarcely, in some posterior cells. At 90h, most cells contained N-Cadherin at various levels, whereas E-Cadherin repression was visible only in a small region. At 96h and 102h, a restricted cellular domain only retained high expression of E-Cadherin. These results suggest that a stepwise process underlies this transition, with an initial widespread activation of N-Cadherin, followed by a progressive and region-specific repression of E-Cadherin, the latter repression being the rate limiting factor of this transition.

We evaluated whether this ordered EMT was associated with the acquisition of specific cellular identity by using either SOX2 or TBX6 to identify NMP or mesodermal cells, respectively (Fig. 2E, F, Fig. S4B, C). While TBX6 was not detected at 78h, SOX2 was transcribed at various levels, with a salt-and-pepper distribution. At 90h, TBX6 and SOX2 expression domains were poorly organised; TBX6-positive cells were often, though not always at the periphery, and most of them displayed high N-Cadherin levels. Instead, SOX2 was frequently excluded from the periphery and showed inconsistent expression at the centre. At 96h and 102h, however, this somewhat unclear situation was resolved and the expression domain were well established with mesodermal cells showing low levels of E-Cadherin expression and high N-Cadherin content. Cells with high E-Cadherin levels also expressed SOX2. Finally, at 114h and 120h, the E-Cadherin level was low throughout, including the mesodermal and neuronal/NMP cells. These results support self-organization as largely arising from a cell-sorting mechanism provided by the stepwise, cell type specific timing of a E-to N-Cadherin switch occurring in both mesodermal and neuronal lineages, yet at a distinct time. our analysis suggests that a difference in the timing of E-Cadherin repression between the NMP population and the mesodermal cells may be involved in the antero-posterior axis formation.

### Timing of cell fate transition

To accurately document the timing of cell fate transitions at the transcriptional level, we combined the single cell RNA-seq datasets with our comprehensive transcriptome time-series. We assessed the dynamic expression of the top ten markers extracted from each cellular cluster, between 48h and 96h, with a 6h resolution (Fig. S3D). Gastruloid cells generally clustered according to their developmental timing, with a major cluster including the 48h up to 84h early time-points, and another cluster comprising late samples (from 72h to 96h). For some time-points, cells were present in both clusters, indicating that the temporal resolution adequately documents the major transitions. This also indicated that, after Chiron treatment, gastruloids are not always perfectly synchronised, as previously reported^14^.

This analysis revealed that neuronal and somitic identities are initiated at the transcriptional level between 78h and 84h. In contrast, repression of the pluripotency program (ESCs, epiblast and pluripotent clusters) started at 72h and was completed by 84h. Finally, transitional states (primitive streak, anterior primitive streak) are initiated as early as 60h and peak at 72h. Therefore, the initiation of the primitive streak identity occurred before the repression of the pluripotency program, reminiscent of the global N-Cadherin expression prior to E-Cadherin repression. These results suggest that the initiation of a new transcription program precedes the termination of the program implemented before.

### Effects of N-Cadherin inactivation

Since these results identified a link between a coordinated cadherin switch and cell specification during gastruloids development, we set out to assess the role of this cadherin transition. First, we inactivated N-Cadherin by deleting the *Cdh2* promoter sequences as well as the first exon and produced mutant gastruloids (Fig. 3A). The elongation of *Cdh2*^-/-^ gastruloids was unexpectedly enhanced, along a single axis (Fig. 3B), with high reproducibility (Fig. S5A). Posterior markers like *Hoxb9*, T/BRA or TBX6 were properly polarised in these mutant specimens supporting a largely normal organization of the posterior domains (Fig. 3C-D, Fig. S5B). Unlike in control gastruloids, alternating high *versus* low intensity segmental expression of *Meox1* was observed (Fig. 3E). In addition, FOXC1 was detected in numerous segments surrounding a FOXC1 negative tubular structure (Fig. 3F), reminiscent of patterned embryonic somites surrounding a neural tube, a situation never observed in native gastruloids (Fig. S5C).

**Fig. 3:**
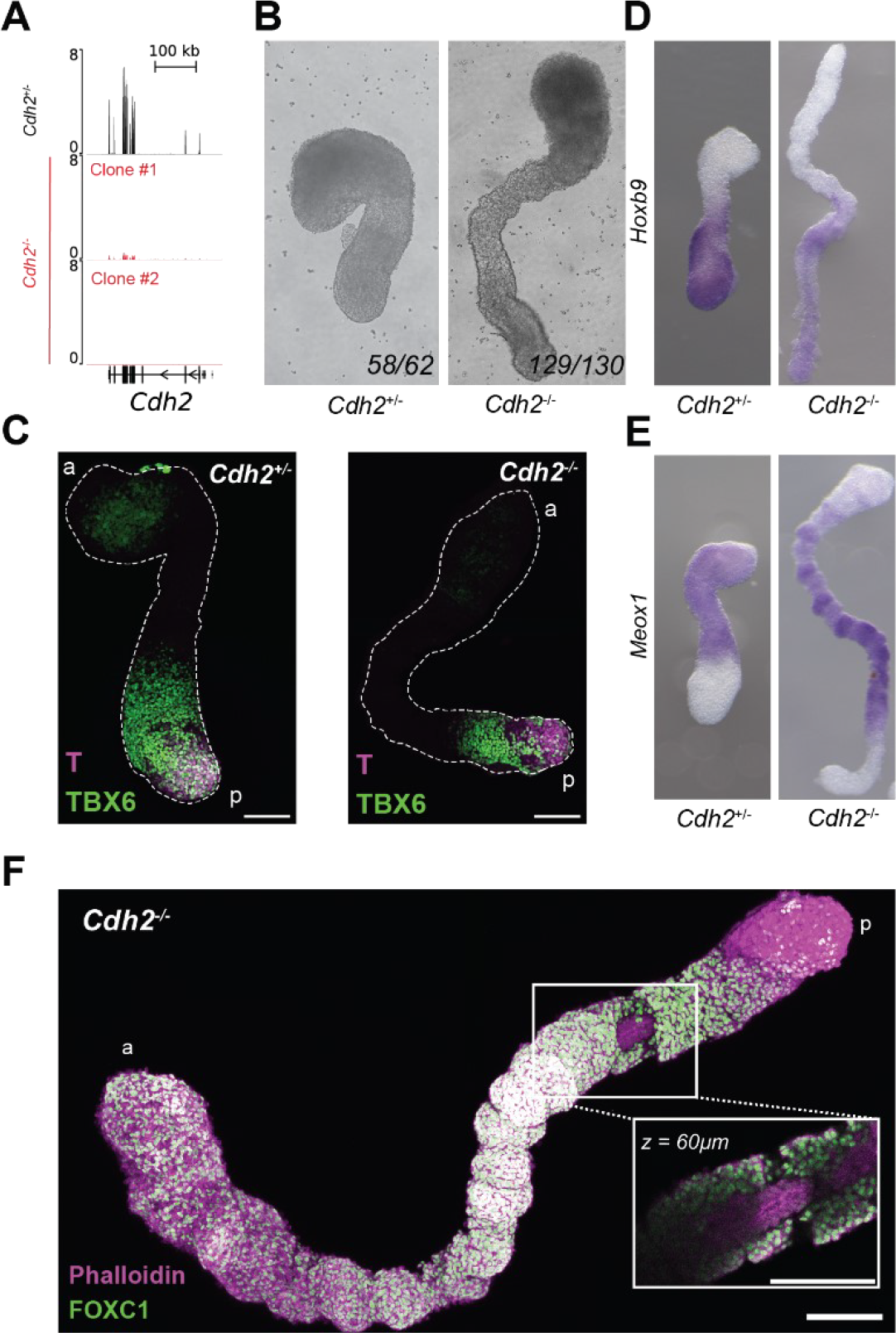
N-Cadherin inactivation releases gastruloid morphogenetic potential. **(A)** Normalised coverage of bulk RNA-seq (read per million) at the *Cdh2* locus in *Cdh2*^+/-^ and two clones of *Cdh2*^-/-^ in 120h gastruloids, average of two replicates. The deleted *Cdh2* promoter region and exon 1 is highlighted in yellow. **(B)** Morphology of 120h gastruloids derived from *Cdh2*^*+/-*^ and *Cdh2*^*-/-*^ ESCs seen with brightfield imaging. **(C)** Maximum intensity projection of immuno-fluorescence against TBX6 (green) and T/Bra (magenta) on *Cdh2*^*+/-*^ and *Cdh2*^*-/-*^ 120h gastruloids. Scale bars are 100 μm, at least 4 gastruloids were imaged on a confocal microscope Sp8 (Leica). For each condition, a designate the anterior side and p designate the posterior side. Gastruloids contours are represented by the dashed line. **(D, E)** *In situ* hybridisation for *Hoxb9* **(D)** and *Meox1* **(E)** on 120h gastruloids derived from *Cdh2*^*+/-*^ (left) and *Cdh2*^*-/-*^ (right) ESCs. **(F)** Maximum intensity projection of immuno-fluorescence for FOXC1 and F-Actin (Phalloidin) in *Cdh2*^-/-^ (KO) 120h gastruloids. The square indicates the area used for blow up of a single z section. All scale bars are 100 μm.

Next, we compared the transcriptome of 120h *Cdh2*^-/-^ gastruloids to those of their *Cdh2*^-/+^ and *Cdh2*^+/+^ controls and, in marked contrast with the strong mutant phenotype, few if any differences in transcripts were scored *Cdh2* expression being the only significant change (Fig. S5D). Patterned somites surrounding a neural tube were only observed when gastruloids were embedded into Matrigel at 96h^17,18^, whereas non-embedded counterparts never displayed any somite-like structure. Therefore, we propose that trunk-like morphogenesis observed in ECM-driven gastruloids arises from the observed segregation of N-Cadherin^18^. The absence of alterations in the mutant *Cdh2*^-/-^ transcriptome illustrates well the decoupling that may exist between morphogenetic events, on the one hand, and the genetic control of those cells, on the other hand. In this view, we can exclude a role for *Cdh2* in establishing the antero-posterior axis through a cell sorting mechanism. Instead, *Cdh2* inactivation uncovered a self-organizing potential leading to a higher resemblance with embryonic structure.

### Snai1 and the robustness of self-organization

During gastrulation, *Snai1* is essential to allow proper mesodermal cell migration by inducing an EMT in the primitive streak and repressing E-Cadherin^29^. In gastruloids, *Snai1* is expressed first in the nascent mesodermal population at 72h and subsequently, in presomitic mesoderm at 96h and 120h (Fig. S6A). Since cells expressing *Snai1* at 72h are also the first to initiate repression of E-Cadherin (Fig. 2C, Fig. S3A), we sought to prevent E-Cadherin repression in mesodermal cells by eliminating its repressor SNAI1 through deletion of both its first exon and promoter (Fig. 4A). At 96h, most *Snai1*^-/-^ mutant gastruloids were spherical while wild type counterparts had frequently broken symmetry and initiated elongation (Fig. S6B). At 120h, the self-organization capacity of mutant gastruloids was impaired (Fig. 4B, Fig. S6C) leading to variable types of structures. Some *Snai1*^-/-^ gastruloids remain round or ovoid (42%), others were composed of multiple buds emanating from a central portion (38%) and 20% formed structures with a single elongating axis. This variability was seen consistently, yet the relative proportions of phenotypes were batch-specific.

**Fig. 4:**
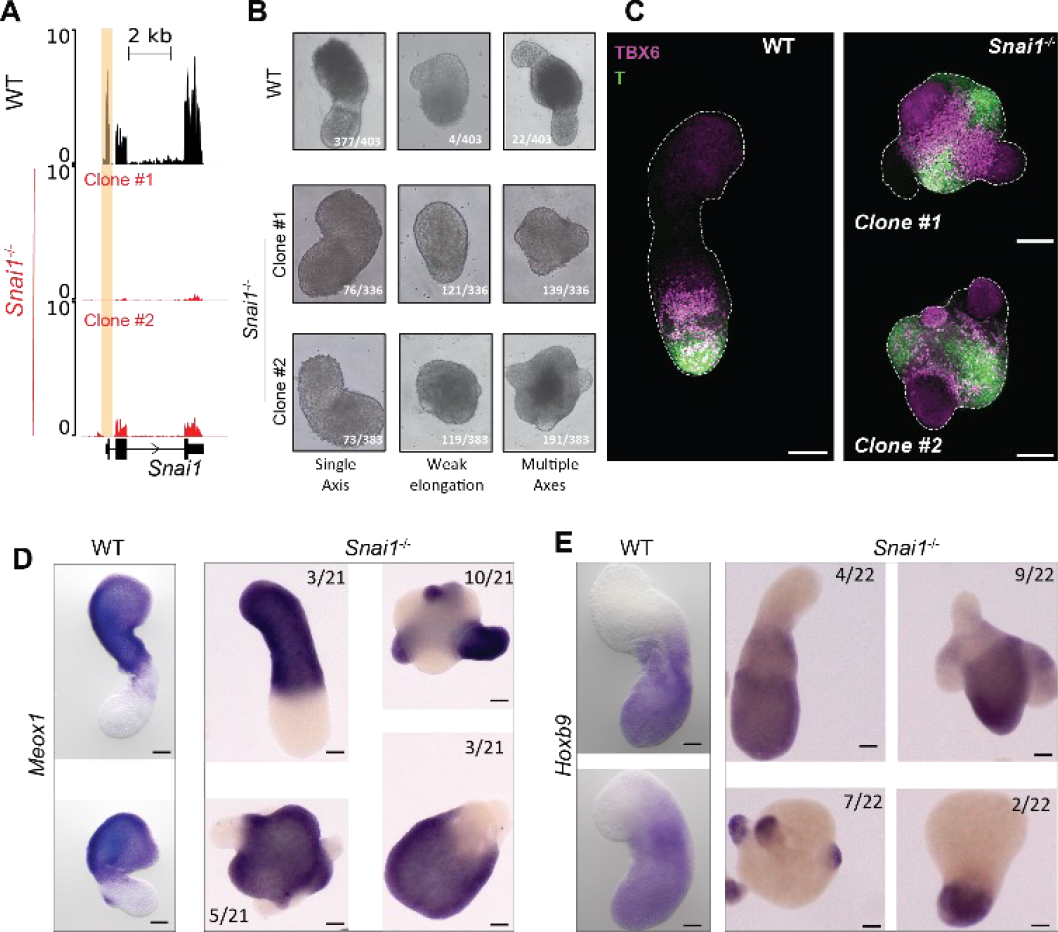
Snai1-mediated repression of E-Cadherin potentiates self-organization. **(A)** Normalised coverage of bulk RNA-seq (read per million) at the *Snai1* locus in WT and *Snai1*^-/-^ 96h gastruloids (average of two replicates). *Snai1* promoter region and exon 1 is highlighted in yellow. **(B)** Brightfield imaging showing the morphology of 120h gastruloids derived from *Snai1*^+/+^ and two clones of *Snai1*^-/-^ ESCs. Results are derived from seven experimental batches **(C)** Maximum intensity projection of immuno-fluorescence for TBX6 (Magenta) and T/Bra (Green) on WT and *Snai1*^-/-^ 120h gastruloids. Scale bars are 100 μm, at least 4 gastruloids were imaged on a confocal microscope Sp8 (Leica). Gastruloids contours are represented by the dashed line. **(D-E)** *In situ* hybridisation for *Hoxb9* **(D)** and *Meox1* **(E)** on 120h gastruloids derived from wild type (left) and *Snai1*^-/-^ (right) ESCs. Four different phenotypes of *Snai1*^-/-^ are observed and represented with their respective proportions. All scale bars are 100 μm.

In addition, *Snai1*^-/-^ gastruloids failed to produce a single posterior pole of T/Bra positive cells. Instead, they frequently formed two posterior poles (Fig. 4C), which tended to be linked by a bridge of E-Cadherin-positive cells (Fig. S6D). In the 120h ovoid or weakly elongated mutant gastruloids, TBX6 expression was limited, whereas multiple axes-containing specimens expressed TBX6 both centrally and at the bases of the protrusions (Fig. S6E). At this stage, E-Cadherin was still detected, indicating that *Snail1*^-/-^ gastruloids had failed to properly downregulate E-Cadherin. In such 120h mutant gastruloids with multiple axes, *Meox1* was expressed sometimes within protrusions, sometimes in the central part or sometimes in the anterior portion where it should normally be transcribed (Fig. 4D). Posterior markers like *Hoxb9* showed a similar variation in transcripts distribution (Fig. 4E). We concluded that *Snai1* plays an essential role during self-organization, in particular by securing the robustness of the process.

### *Snai1* and the repressive tempo of pluripotency

Finally, we investigated how *Snai1* impacts self-organization and pattern formation by initially focusing on both E- and N-Cadherin dynamics in *Snai1*^*-/-*^ gastruloids. If anything, N-Cadherin expression was only weakly impacted in *Snai1*^-/-^ mutants and expression was still detected at 90h (Fig. 5A, Fig S7A). In contrast, the timing of E-Cadherin repression was severely affected: while control gastruloids showed a regionalised loss of E-Cadherin as early as 90h and consistently at 96h, *Snai1*^*-/-*^ specimen expressed high levels of E-Cadherin at both 96h and 102h (Fig. 5A). Initiation of repression was only visible at 114h and 120h. These results suggest that *Snai1* triggers a rapid E-Cadherin repression in mesodermal gastruloid cells. Instead, NMPs and neuronal cells undergo a slower E-Cadherin loss until around 120h. In the absence of *Snai1*, neuronal and mesodermal cells behave more synchronously, undergoing slow and progressive decrease in E-Cadherin expression. In addition, SOX2 expression, which is normally restricted to a posterior NMP portion (Fig. S7B), was also impacted by *Snai1* and was maintained throughout a large portion of the gastruloid until 102h (Fig. 5B). At both 114h and 120h, when *Snai1*^*-/-*^ gastruloids formed protrusions, SOX2 became restricted to the central body, suggesting that such cells may either correspond to uncommitted NMPs or, alternatively, that they may have kept features of pluripotency.

**Fig. 5:**
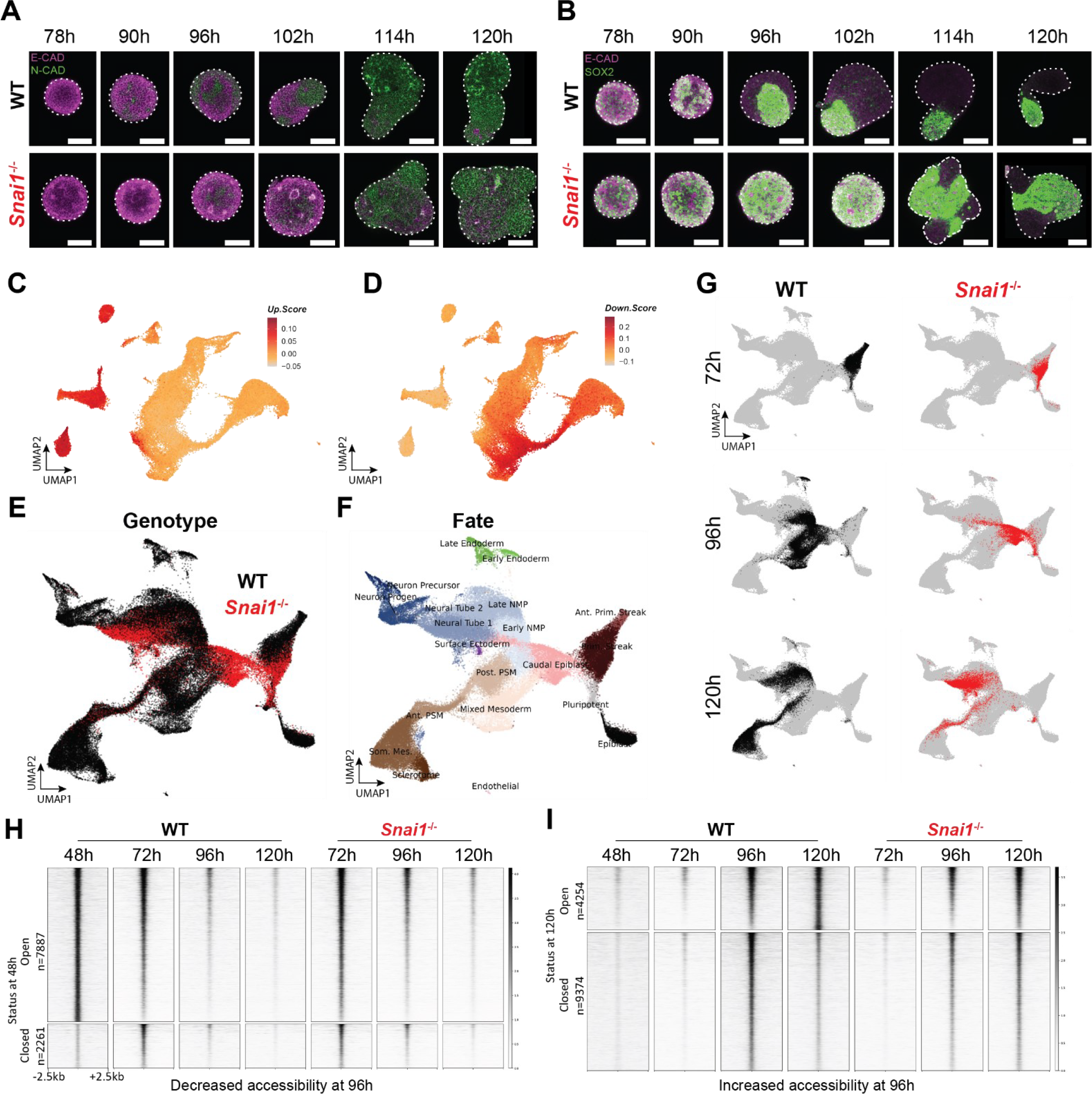
Snai1 controls the repressive tempo of an early pluripotency associated program. **(A-B)** Immuno-fluorescence for E-Cadherin (Magenta) and N-Cadherin (green) **(A)**, or E-Cadherin (Magenta) and SOX2 (Green) **(B)**, on WT (top) and *Snai1*^-/-^ gastruloids (bottom) at the indicated time points. Gastruloids contours are represented by the dashed line. z-stack is 20 μm in the gastruloids. At least 5 gastruloids were imaged per condition. Imaging was done with 20x magnification on a confocal microscope Sp8 (Leica). All compared conditions had the same illumination and acquisition settings. Scale bars are 100 μm. **(C-D)** UMAP plot of single cell RNA-seq experiment from wild type gastruloids (from **Fig 1A, B)**, coloured based on Gene Set Enrichment Analysis (from **Fig. S8A**) of the list of upregulated **(C)**, or downregulated **(D)** genes in *Snai1*^-/-^ compared to wild type 96h gastruloids cells **(E-F)** UMAP plot of single cell RNA-seq of ∼72’000 cells from WT and *Snai1*^-/-^ gastruloids at 72h, 96h and 120h. coloured by **(E)** Genotype, wild type (black) or *Snai1*^-/-^ (red) or by clusters **(F)**. Of note the 48h, 144h and 168h are also displayed, but only analysed from a wild type context. Prim. Streak, Primitive Streak; NMPs, Neuro-Mesodermal Progenitors; Progen., Progenitor; Post./Ant. PSM, Posterior/Anterior Pre-Somitic Mesoderm; Som. Mes, Somitic Mesoderm.(**G**)UMAP plot highlighting from each timepoint, wild type cells (black, left panels) and *Snai1*^-/-^ (red, right panels), while the remaining cells are in grey. **(H-I)** Heatmap showing ATAC-seq signal from wild type and *Snai1*^-/-^ gastruloids at 72h, 96h and 120h at sites losing **(H)** or gaining **(I)** accessibility at 96h compared to 72h. Of note, the 48h condition was only performed on wild type gastruloids. Sites with differential accessibility were defined with abs(log2FC) < 1, padj< 0.05). All values are normalised by the number of reads in peaks, peaks are centred and ranked by ATAC-seq signal at 72h (**H**) or 96h (**I**) from wild type gastruloids. Status at 48h **(H)**, or 120h **(I)** was determined by the overlapping with a confident peak (see Methods). WT 48h is the average of two replicates and KO 72h and 96h is the average of two clones.

To identify the early-responding targets of *Snail1, w*e analysed *Snail1* mutant transcriptomes at 96h, i.e., when the abnormal phenotype starts to be observed. Consistent with its known repressor activity, transcriptional up-regulation was the most frequent change (862 up-regulated genes *versus* 432 down-regulated), with *Cdh1* (E-Cadherin) being the most significant target (Fig. S8A). Amongst other mis-regulated genes, *Tbx6, Aldh1a2* and *Pcdh19*, all part of the mesodermal program, were down-regulated, suggesting a role for *Snail1* in securing mesodermal fate. To infer which genetic program is impacted by the loss of *Snai1*, we generated an enrichment score of down-regulated *versus* up-regulated genes in our single cell time-course analysis (Fig. 1A). The up-regulated program was enriched in all the ‘early’ cell populations such as ESCs, epiblast, pluripotent and primitive streak (Fig. 5C, Fig. S8B) supporting a requirement of *Snai1* to repress an early transcriptional program, already active in ESCs and only lost at ca. 96h gastruloids, once cells have committed to either the NMPs/neuronal, or the mesodermal lineages. In contrast, genes thar are down-regulated at 96h strongly associated with mesodermal lineages, in particular somitic mesoderm, as well as the neuronal/NMP lineages, to some extent (Fig. 5D, Fig. S8B).

To further evaluate the potential cell-specific defects occurring in mutant gastruloids, we performed a single cell transcriptomic time-course comparing control and *Snai1*^*-/-*^ specimens at 72h, 96h and 120h, (Fig. 5E-G, Fig. S8C-D). *Snai1*-deficient gastruloids lacked a nascent mesoderm population at 72h, while all remaining cells had activated the primitive streak program, with expression of *T, Fst* and *Sp8* (Fig. 5E-G, Fig. S8E). At 96h, most *Snai1*^*-/-*^ cells were in a premature transcriptional state similar to the caudal epiblast program (Fig. 5F, Fig. S8D), i.e., failing to express mesodermal marker genes like *Tbx6* or *Msgn1* (Fig. S8F). Instead, the expression of genes normally down-regulated and associated with pluripotency (*Utf1, Pou5f1*) was maintained (Fig. S8G). At 120h, *Snai1*^*-/-*^ gastruloids contained properly specified cells, yet in altered proportions and stages of differentiation (Fig. S8D). Mesodermal cells were strongly reduced in number and their differentiation was delayed, only reaching an anterior PSM state and lacking the expression of somitic markers like *Pax3, Meox2* or *Pax1* (Fig. S8H). In contrast, neuronal differentiation was over-represented (Fig. S8D), progressing even further than expected when compared to controls, as illustrated by numerous cells expressing *Pax6* and *Irx3* (Fig. S8I). Therefore, in addition to failing to form organised structures, gastruloids derived from *Snai1*^-/-^ cells lack the proper temporal control of cell fate progression. Finally, because *Snai1* encodes a DNA-binding transcriptional repressor of both E-Cadherin and a global pluripotency program^30,31^ during development and tumorigenesis^32–34^, we used an ATAC-seq approach in control and *Snai1*^*-/-*^ gastruloids at 72h, 96h and 120h to identify where and when *Snai1* might impact the chromatin accessibility landscape. At 96h, i.e., when the pluripotency-associated transcriptional program was incorrectly maintained in mutants (Fig. S8G), we selected sites with decreased accessibility between 72h and 96h control gastruloids (Fig. 5H). Out of 10’148 such sites identified, 7’887 (ca. 80%) were already opened at 48h, suggesting they were part of an early pluripotency-associated program. In *Snai1*^-/-^ gastruloids, this transition from open to closed sites at 96h was severely impaired, an abnormal situation not even fully resolved at 120h. This was best exemplified by the E-Cadherin (*Cdh1*) and the pluripotency factors *Oct4* (*Pou5f1*) loci (Fig. S9A). In contrast, chromatin sites becoming open at 96h in control gastruloids remained unimpacted by the absence of *Snai1* (Fig. 5I, S9B) despite a failure to activate the corresponding transcriptome (Fig. 5D). We interpret this as a decoupling between the opening of a chromatin landscape, on the one hand, and its following transcriptional activation, on the other hand. It is possible that the repression of a previously deployed program is required to progress to more advanced differentiation states. This would be consistent with the observed delay in the transcriptional onset of both the neuronal and somitic programs in *Snai1* mutant gastruloids, which occurred only in cells where pluripotency was repressed (Fig. S8F-I).

## CONCLUSION

In this work, we used a multi-scale approach to investigate potential mechanisms underlying the outstanding capacity of gastruloids to self-organizing during the first five days of their development. We identified a mechanism leading to the specific sorting out of various cellular population based on the coordination, at the cellular level, of an E-to N-Cadherin switch. We report that N-Cadherin is largely dispensable for self-organization and that its inactivation leads to an improvement to an embryonic-like morphology resembling trunk-like structures. We interpret this as the capacity of N-Cadherin to prevent a further step in self-organization to occur in control gastruloids (i.e., a somitic-like segmentation), an effect that can be overcome by some compounds delivered by the Matrigel. The Cadherin switch is associated with an exit from pluripotency, as evaluated by transcriptomes and chromatin accessibility. This is orchestrated by the transcriptional repressor *Snai1*, which sets the tempo of both this repression and the cell fate transitions, leading to the observed decrease in chromatin accessibility around genes associated with a pluripotency program. We suggest that this chain of events is an integral part of the self-organizing mechanism, which is likely well conserved between genuine embryos and stem cell based *in vitro* models. Similar to normal embryos, this process seems to be very robust and hence it may explain the high reproducibility observed during the production of gastruloids^23,11^.

## METHODS

### mESCs and gastruloids culture

mESCs (129/svev, EmbryoMax© CMTI-1) were cultivated in an humidified incubator (5% CO2, 37°C), between Passage 15 and 35 in 6-well plates (Corning, 354652) using a 2i^35^, LIF DMEM medium composed of DMEM + GlutaMAX (Gibco, 10566-024) supplemented with 10% ES certified FBS (Gibco, 16141079), non-essential amino acids (Gibco, 16141-079), sodium pyruvate (Gibco, 11360-039), beta-mercaptoethanol (Gibco, 31350-010), penicillin/streptomycin (Gibco, 15140-122), 100 ng ml−1 of mouse LIF (custom made by the EPFL Protein Production and Structure Core Facility), 3 μM of GSK3 inhibitor (CHIR99021, FBM-10-1279-5MG) and 1 μM of MEK1/2 inhibitor (PD0325901, Selleckchem, S1036)^12^. Lif, MEKi and GSK3i were aliquoted and kept at −20°C, complete medium, were not kept more than one week. When a density of 200’000 cells per cm^2^ was reached, cells were washed with PBS (Gibco, 10010023) and dissociated using Accutase© (StemPro Ref: A11105-01). They were resuspended in DMEM medium, counted on a Countess 3 automated cell counter (Invitrogen) and reseeded at 200 000 cells or 80 000 cells per well when kept for two or three days respectively.

The differentiation protocol for gastruloids was previously described^10^. Briefly, mESCs were dissociated with Accutase treatment, washed and resuspended in prewarmed N2B27 medium composed of 50% DMEM/F12 (Gibco, 11320033) and 50% Neurobasal (Gibco, 21103049) supplemented with 0.5× N2 (Gibco, 17502048) and 0.5× B27 (Gibco, 17504044). In total, 300 cells were seeded in 40 μl of N2B27 medium in each well of a low-attachment, rounded-bottom 96-well plate (Corning, 7007). 48 hours after aggregation, 150 μl of N2B27 medium supplemented with 3 μM of GSK-3 inhibitor was added to each well. Then, 150 μl of the medium was replaced every 24 h until collection. The timing of GSK-3 inhibitor and gastruloids medium change was always within 1 h of the time of aggregation. Harvesting of gastruloids was performed regardless of the morphology at all analysed stages except when the whole plate of gastruloids failed to elongate, those batches were excluded from the analyses.

### Mutant mESCs generation

Wild-type mouse embryonic stem cells (EmbryoMax 129/svev) were used to generate all the mutant cell lines used following the CRISPR/Cas9 genome editing protocol described in^36^. sgRNA targeting guides (**Supplementary Table 1)** were cloned into a Cas9-T2A-Puromycin expressing plasmid containing the U6-gRNA scaffold (gift of A. Németh; Addgene plasmid, 101039). 8 μg of plasmid was used to transfect mESCs using the Promega FuGENE 6 transfection kit. Those were dissociated 48 h later for puromycin selection (at 1.5 μg/ml). Clone picking was conducted 5–6 days later, and resistant clones were plated on gelatinized 96-well plates (Corning, 354689) and analysed by PCR screen using the MyTaq PCR mix kit (Meridian Bioscience) and specific primers surrounding the targeted region (**Supplementary Table 2**). Deletions or insertions were verified for both alleles by Sanger sequencing (**Supplementary Table 3**). All knock-out lines were obtained directly after one round of transfection. The *Mesp2* reporter line was generated by integrating a p2a-H2BmCherry in frame with the *Mesp2* coding sequence just upstream of the Stop codon. This insertion was done on a hemizygote allele of the *Mesp1/2* locus where one copy of the whole locus was deleted. 8 μg of the template repair plasmid was along with the sgRNA plasmid. Of note, in one of the two *Snai1* clones that was analysed, although the deletion was successfully obtained in both alleles, an alternative promoter was activated leading to a truncated *Snai1* mRNA. However, we could not identify a different phenotype between the two clones, suggesting that this alternative transcript is not functional. One of the *Cdh2* mutant clone has a slight contamination by ∼1% of wild type cells. RNA-seq was performed on two clones, including the one with the contamination. Staining however was only performed in the contaminated clone, but 5 clones were analysed phenotypically by wide-field microscopy and all behave similarly suggesting that this contamination did not impact our analysis in any way.

### Gastruloid dissociation

At the indicated stages, gastruloids were collected as a pool in a 15 ml tube and were washed once with PBS. After centrifugation, the gastruloids pellet was dissociated with 200 μl of Accutase (StemPro) and incubated at 37°C for 3 minutes, full dissociation was achieved by mechanical dissociation (pipetting). Following centrifugation, the cell pellet was then resuspended in the adequate volume of N2B27 medium (200 μl per well) and replated in their original round bottom 96 well plate. The plate was centrifuge for 1 minute at 300g and placed back in the incubator for analysis.

### Whole-mount In Situ Hybridization (WISH)

Gastruloids were collected at the indicated stage and processed following a previously reported WISH procedure^10^. Briefly, they were fixed overnight in 4% PFA at 4°C and stored in methanol at −20°C until ready for processing. Each sample was rehydrated and prepared with Proteinase K (EuroBio) at 2.5 μg/ml for 2 minutes. They were incubated in a blocking solution at 68°C for 4 h before incubation overnight with specific digoxigenin-labelled probes (see **Supplementary Table 4**) at a final concentration of 100-200 ng/ml. The next day, samples were washed and incubated with an anti-DIG antibody coupled to alkaline phosphatase (Roche, 1:3,000). Staining used BM-Purple (Roche).

### Whole-mount Immuno-Fluorescence

Gastruloids were collected, washed with PBS and then fixed in 4% PFA; 1% BSA for 1 h in Eppendorf tubes at room temperature on a roller. Then, they were washed three times in PBS for 5 minutes, and kept at 4°C until further processing. Then, permeabilization was achieved by 3 x 10 minutes incubation in 0.2% Triton-X/PBS; 10%FBS (PBS-FT). Then they were blocked PBSFT for one hour at 4°C. Samples were then incubated with the primary antibodies in PBSFT for 48 hours at 4°C. Then, a series of two 5-minutes, three 15-minutes and four 1-hour washes were done in PBSFT. Then incubation with the secondary antibody, and other fluorescent reagents (DAPI, Phalloidin) was performed overnight at 4°C. Stained gastruloids were washed with three 20-minutes washes in PBSFT and were mounted on slides, PBSFT was removed and Refractive index matching solution (2.5M Fructose, 60% Glycerol, 40% water) was then added to cover the stained gastruloids which were imaged at least 24 hours after mounting. The list of primary and secondary antibodies is in **Supplementary Table 5**.

### Image acquisition

Pictures of *in situ* hybridizations were done with an Olympus DP74 camera mounted on an Olympus MVX10 microscope using the Olympus CellSens Standard 2.1 software. Immunofluorescence were imaged on a point scanning confocal microscope SP8 upright with a Lumencor Sola SM II LED laser, Hybrid detectors (HyD) and a HC PL APO 20x objective. All confocal samples that are directly compared were imaged with the same acquisition settings (laser intensity, resolution, detector gain…). Widefield live imaging was performed on an IncuCyte S3 (Sartorius) using the Spheroid module and if required fluorescence acquisition used the default parameters (400ms exposure time for the red channel).

### Image analysis

Gastruloids classification for dissociation experiment, or phenotypical analysis of *Snai1* mutant was performed manually. Image analysis was done using ImageJ software analysis. For membrane quantification of E- and N-Cadherin signal, we first built a synthetic membrane channel by adding the E- and N-Cadherin channels. A pixel classifier (ilastik^37^) was trained to define membrane ROI from the sum of E-Cadherin and N-Cadherin signals. Then, on each individual channel, a median filter of 1 μm radius was applied and a modified version JACOP^38^ script was used to quantify the expression of E and N-Cadherin at membrane ROI and produce the fluorogram shown in **Fig. S3A**.

### NGS analysis

All NGS CPU-demanding analyses were computed using a local galaxy server^39^. All coverage plots were realised with pyGenomeTracks v3.8^40,41^ using the facilities of the Scientific IT and Application Support Center of EPFL.

### Bulk RNA-seq

Gastruloids were harvested and pooled in a 2 ml Eppendorf tube at the indicated time-points. After medium removal, gastruloids were washed once with PBS, pelleted and stored at −80 °C until RNA extraction. RNeasy Mini kit (Qiagen) with on-column DNase digestion was used for RNA extraction following manufacturer’s instructions. RNA quality was assessed on a TapeStation TS4200, all RNA samples showed quality number (RIN) above 9.5. RNA-seq library preparation with Poly-A selection was performed with 1000 ng (for time-course analysis) or 500 ng (for mutant gastruloid analysis) of RNA using the Illumina stranded mRNA ligation and following the manufacturer’s protocol 1000000124518 v01. Library quality was assessed with a Libraries were quantified by qubit DNA HS and profile analysis was done on TapeStation TS4200. Libraries were sequenced on HiSeq 4000 Illumina, with paired end 75 bp reads. For the gastruloid time-course, three independent batches of gastruloids were analysed for each time-points. For *Cdh2* and *Snai1* mutant gastruloid analysis, two knockout clones were analysed each in two independent batches.

### Bulk RNA-seq Analysis

Raw RNA-seq reads generated in this study were trimmed to remove Nextera adapters or bad quality bases (Cutadapt v4.0^42^ -a CTGTCTCTTATACACATCTCCGAGCCCACGAGAC -A CTGTCTCTTATACACATCTGACGCTGCCGACGA -q 30 -m 15). Mapping and counting were performed on filtered reads on the mouse genome mm10 with STAR v2.7.8a^43^ with a custom gtf based on Ensembl v102^44^ and ENCODE parameters. FPKM values were obtained by Cufflinks v2.2.1^45,46^ with options --no-effective-length-correction -b ‘mm10.fa’ --multiread-correct --library-type fr-firststrand --mask-file ‘chrM_mm10.gtf’ --max-bundle-length 10000000 --max-bundle-frags 1000000 (where chrM_mm10.gtf contains a transcript on each strand of the whole chrM). Uniquely mapped reads were filtered with bamFilter^47^ v2.5.1. Coverages were computed with BEDTools^48^ v2.30.0 for each strand using the ‘-scale’ parameter to normalize to million uniquely mapped reads. Average between duplicates was performed with bigwigAverage v3.5.4 from deepTools^49^. PCA, correlation matrices and clustering were performed on log2(1 + FPKM) values of the 2000 most variant genes. Visualization was done in R v4.2.1, Heatmaps were generated with the Pheatmap package (v1.0.12) using log2(1 + FPKM) values. The sample correlation matrix was obtained using the 2000 most variable genes using the “ward.D2” clustering method on the spearman inter-sample correlations. The heatmap on single-cell marker genes was computed scaling the log2(1+FPKM) values per gene. The sample clustering was done using Pearson’s correlation from the visualised genes with the “ward.D2” method. Differential gene expression between wild type and the different mutant lines was done using DESeq2^50^ v1.42.0. All samples were of good quality and none were excluded from the analysis. Genes were called differentially expressed when they had both abs(log2fc) > 1 and p-adj < 0.05.

### Reanalysis of Bulk RNA-seq from GSE106227

Single-reads fastqs were retrieved from the SRA server using fasterq-dump v3.0.5. They were trimmed for bad quality and Truseq adapters with Cutadapt v4.0 (-a CTGTCTCTTATACACATCTCCGAGCCCACGAGAC -q 30 -m 15). Mapping and FPKM calculations were performed as described above. Coverage was obtained using only uniquely mapped reads. The heatmap on markers for Epiblast, Pluripotent, PS and APS was computed on Pearson’s correlation on log(1 + FPKM) values. The clustering was done with euclidean distance and ward.D2 method.

### Single-cell RNA-seq

Gastruloids were collected, pooled in a 1.7 ml Eppendorf tube and washed twice in 1 ml of PBS. The number of gastruloids used in each condition was chosen to ensure that more than 300 000 cells were obtained. They were then dissociated in 100 μl of Accutase (Stempro) for 5 minutes at 37°C. Full dissociation was completed using mechanical dissociation and verified to ensure the absence of doublet cells. Cells were then resuspended in 200 μl of PBS and counted on a Countess 3 automated cell counter (Invitrogen). All centrifugations were done at 400g for 5 minutes. Some samples were then analysed with the standard 10x genomics pipeline (Chemistry v3 and v3.1), while others also included a CellPlex procedure to multiplex samples (see **Supplementary Table 6** for list of samples and the method used). For the sample treated with the normal 10x procedure, after counting, dissociated cells were diluted in PBS BSA 0.04% (Millipore, A1595) to 1×10^6^ cells/ml. For samples treated with the CellPlex multiplexing approach, cells were pelleted and incubated for 5 minutes in 50 μl of cell multiplexing oligos (3’ CellPlex Kit Set A, PN-1000261) at room temperature. They were then washed three times with 1 ml of PBS 1% BSA and finally they were resuspended in 110 μl of PBS 1% BSA and counted. The multiplexed samples were then pooled in equal proportions for each targeted sample. Both CellPlex and standard 10x Genomics sample were then filtered using a 40 μm cell strainer (Flowmi, BAH136800040). The pool of cells was counted and subjected to single cell RNA-seq following manufacturer’s recommendation similarly as non-CellPlex samples (protocol CG000388 Rev A). For standard 10x samples, we aimed for a recovery of ca. 6000 cells per sample while for multiplexed single cell RNA-seq we aimed for a recovery of maximum 24 000 cells as more doublet can be resolved with this method. cDNA preparations were performed according to 10x Genomics recommendations, amplified for 10–12 cycles and verified on fragment analyser. Both cell multiplexing oligo, and gene expression libraries were sequenced on a HiSeq 4000 (ref 15065681 v03) or a Novaseq (Illumina protocol #1000000106351 v03) with the cbot2 chemistry.

### Single-cell RNA-seq analysis

Gene expression libraries were analysed similarly for standard 10x and CellPlex samples to generate the cell-gene expression matrices. For both Mapping, demultiplexing of cell barcode and quantification was performed using rna_STARsolov2.7.10b^51^ on the mm10 reference genome, and a modified gtf file based on Ensembl v102 (doi: zenodo.org/doi/10.5281/zenodo.10079672), with the parameters: --soloBarcodeReadLength 1 --soloCBstart 1 --soloCBlen 16 --soloUMIstart 17 --soloUMIlen 12 -- soloStrand Forward --soloFeatures Gene -- soloUMIdedup 1MM_CR --soloUMIfiltering - -- quantMode TranscriptomeSAM GeneCounts -- outSAMattributes NH HI AS nM GX GN CB UB -- outSAMtype BAM SortedByCoordinate --soloCellFilter None --soloOutFormatFeaturesGeneField3 ‘Gene Expression’ --outSAMunmapped None -- outSAMmapqUnique 60 --limitOutSJoneRead 1000 -- limitOutSJcollapsed 1000000 --limitSjdbInsertNsj 1000000. Then DropletUtils^52,53^ v1.10.0 was used to remove barcodes which were not corresponding to cells (Method EmptyDrops, lower-bound threshold = 100, FDR threshold = 0.01). Cell multiplexing oligo (CMO) libraries which identify the sample of origin in multiplexed experiments were quantified using CITE-seq-Count v1.4.4^54^ --cell_barcode_first_base 1 -- cell_barcode_last_base 16 --umi_first_base 17 -- umi_last_base 28 --bc_collapsing_dist 1 -- umi_collapsing_dist 2 --expected_cells 24000 --whitelist ‘3M-february-2018.txt’ --max-error 2. Individual 10x gene expression libraries, were analysed using Seurat v4.3^55^. We first filtered out barcodes with less than 200 identified gene and genes identified in less than three cells. For CMO libraries, the demultiplexing by barcode was done in R using counts from CITE-seq-Counts. Cell barcodes with less than 5 CMO UMI and absent in the Seurat object were discarded. Then sample attribution was performed using demuxmix^56^ using the total number of UMI per cell. Cells which were not classified by demuxmix as singlet (negative, unsure, doublet) were excluded from the analysis. Low quality cells and potential doublets were removed by computing the mean UMI content and the percentage of mitochondrial genes and filtering out barcodes with less than 0.4 times the mean UMI or more than 2.5 times the mean UMI. Only barcodes between 0.05 and 8 percent of mitochondrial UMI were kept. Matrices were normalised and the cell cycle score (using the 2019 updated gene list from Seurat) from these filtered libraries were computed. Then we merged the different samples using the merge command by Seurat. The combined Seurat object was then normalised, 2000 variable features were identified and the data was scaled and regressed by cell cycle score, and percentage of mitochondrial reads. Principal component were then computed using variable genes falling within the 5^th^ and 80^th^ percentile of expression to limit batch effect as performed here^57^. UMAP and k-nearest neighbours were then computed using 30 principal components. The resolution to cluster the data was chosen by ensuring that no cluster was missed, and that no duplicated clusters were found. Cluster annotation was performed manually. Marker identification was done using a maximum of 400 cells per identity to limit the impact of widely variable cluster sizes in the ability to identify markers, these also required a difference of 20% in the proportion of expressing cells and a logFC >0.5. For RNA velocity analysis, Velocyto v0.17.17^58^ was used with the run10x command on the BAM files generated by STARsolo. Then, velocyto.R v0.6 was used to identify the velocity using the RunVelocity function with the settings: spliced.average = 0.2, unspliced.average = 0.02, deltaT = 1, kCells = 20, fit.quantile = 0.02. RNA velocity was then displayed on the UMAP embedding with the show.velocity.on.embedding.cor function with the setting: n = 300, scale=‘sqrt’, cell.colors = ac(subset.seurat$FateColor, alpha=0.5), cex=0.8, arrow.scale = 5, show.grid.flow=TRUE, min.grid.cell.mass = 0.5, grid.n=40, arrow.lwd = 1, do.par = F, cell.border.alpha = 0. In order to study the correlation between *Mesp1* and *Lhx1* genes with other markers, cells were classified in four different categories. Cells with normalised expression at 0 were classified as non-detected (ND), between 0 and 1 as low, 1 and 2 as medium and above 2 as high.

For cross comparison with bulk RNA-seq of *Snai1*^-/-^ at 96h, differentially expressed genes with abs(average of log2FC for both clones) > 2 and adjusted p-value < 0.00005 in both clones were further filtered according to their single cell expression (average exp >0.3 in at least on cluster) were used for scoring and heatmap. Scoring of cells based on this list of genes up-/down-regulated in Snai1^-/-^ was performed with the addModuleScore by Seurat.

### ATAC-seq

Assay for Transposase Accessible Chromatin (ATAC) was performed as described^59^. Gastruloids were collected, pooled in a 1.7 ml Eppendorf tube and washed twice in 1 ml of PBS. They were then dissociated in 100 μl of Accutase (Stempro) for 5 min at 37°C. Full dissociation was completed by mechanical dissociation and verified to ensure proper dissociation. Dissociated cells were then washed in 500 μl of PBS (centrifugation 5 min at 4°C, at a speed of 400g) and resuspended in 200 μl of resuspension buffer RSB (10 mM Tris-HCl pH7.4, 10 mM NaCl, 3 mM MgCl2). 50 000 cells were extracted, pelleted (centrifugation 5 min at 4°C, at a speed of 400g) and incubated 3 minutes on ice in cold lysis buffer (RSB, 0.1% NP40, 0.1% Tween-20, 0.01% Digitonin). Nuclei were then washed in 1 ml of lysis washout buffer (RSB, 0.1% Tween-20) and pelleted at 500g for 10 minutes at 4°C. Pelleted nuclei were resuspended in 50 μl of cold transposition buffer (TD Buffer, 100 nM Tn5 (Illumina, 20034197), 20% PBS, 0.01% Digitonin, 0.1% Tween-20, 10% water) and incubated 30 minutes at 37°C. Each reaction was purified using the Qiagen MinElute PCR purification kit. The reaction was then amplified by PCR using the NEBNext Master Mix and Nextera primers. All libraries were analysed on a fragment analyser before sequencing on a Nextseq 500 with paired-end 75 bp reads.

### ATAC-seq analysis

Raw ATAC-seq reads were trimmed to remove both Nextera adapters and bad quality called bases (Cutadapt v4.0^42^ -a CTGTCTCTTATACACATCTCCGAGCCCACGAGAC - A CTGTCTCTTATACACATCTGACGCTGCCGACGA -q 30 -m 15). Mapping was performed on filtered reads on the mouse genome mm10 with Bowtie2 v2.5.0^43^ (--very-sensitive 2). Resulting Bam files were sorted using samtools. Unconcordant reads or reads mapping to the mitochondrial chromosome were discarded with bamFilter v2.5.2^47^. Duplicates were removed with Picard MarkDuplicates v2.18.2. BAM was converted to BED with BEDTools v2.30.0 in order to use both mates to call peaks. Peaks were called using macs2_callpeak v2.2.7.1. with the options (--gsize ‘1870000000’ --call-summits -- keep-dup ‘all’ --nomodel --extsize ‘200’ --shift ‘-100’ -- bdg). Coverages were converted to bigwig and normalised to the number of million reads mapped less than 500bp from a peak summit. Normalised coverages from wild-type replicates 1 and 2 were averaged with bigwigAverage v3.5.4 from deepTools^49^ as well as the coverages from the two KO clones at 72 and 96h. In order to build a comprehensive and confident list of peaks from the wild-type time-course, at each time-point the following strategy was adopted. The BED with the largest number of reads was down sampled to match the number of reads of the other replicate. Peaks were called using a merged of equally sized BED. Only peaks whose summit was overlapping a peak in each of both replicates were kept. The four lists of confident peaks (one per time-point) were concatenated and merged with BEDTools v2.30.0. The number of reads falling into this comprehensive confident list were quantified with deeptools multiBamSummary v3.5.1^49^. DESeq2 was used to compare the wild-type duplicates of 72h with the wild-type duplicates of 96h to determine the loci with a significant change in accessibility (abs(log2FC) > 1 an adjusted p-value < 0.05). These loci were sorted by average normalised coverage in 72h for the loci with decreased accessibility at 96h and in 96h for loci with increased accessibility at 96h. The status at 48h and 120h was determined as overlapping a confident peak at this time-point. The heatmap was computed with computeMatrix and plotHeatmap from deepTools v3.5.1 on a 5 kb region surrounding the peaks centre. The Snai1 mutant ATAC-seq were only used for visualisation as heatmaps and genome browser views, the list of regions and measure of accessibility changes in time only included wild type samples.

## DATA ACCESSIBILITY

All the sequencing data found in this study can be accessed on GSE247511.

## CODE ACCESSIBILITY

All the code required to reproduced the analysis (imaging, single cell RNA-seq, Bulk RNA-seq and ATAC-seq) can be found on: https://github.com/MayranA/allScriptsFromMayranEtAl2023 (https://doi.org/10.5281/zenodo.10118923).

## MATERIAL ACCESSIBILITY

All the cell lines generated in this project are available upon request.

## CONTRIBUTION

A.M. and D.D. conceived the study and designed the experiments with input from P.O. A.C. and D.K generated the mutant ESC lines. A.M., D.K., M.L., T.L., Y.R., A.C., with help from A.R.A. performed the experiments. A.M. and L.L.D analysed the data. A.M. interpreted the data with input from D.D. and P.O.; A.M and D.D. wrote the manuscript with input from L.L.D.

## ACKNOWLEDGEMENTS

We thank all the colleagues from the Duboule and Andrey laboratories for discussions, Can Aztekin and Anaïs Le Nabec for critical reading of the manuscript. This work was supported in part using the resources and services for sequencing of the Gene Expression Research Core Facility (GECF) at the School of Life Sciences of EPFL, and the BioImaging and Optics Platform of EPFL, in particular, Nicolas Chiaruttini for help in image acquisition and their analysis. In addition, some computational work was performed using the SCITAS facilities of the Scientific IT and Application Support Center of EPFL. This work was supported by funds from the École Polytechnique Fédérale de Lausanne (EPFL, Lausanne), the European Research Council grant RegulHox (588029 to D.D.), the Swiss National Science Foundation (CRSII5_189956, 310030B_138662 to D.D., and 407940_206405 to A.M) and the Human Frontier Science Program (LT000032/2019-L and the Science4Science initiative to A.M.).

