## Supplementary Figure for "Cadherins modulate the self-organizing potential of gastruloids"

### SUPPLEMENTARY MATERIALS

**Fig. S1-9**

**Supplementary Note 1**

**Legend to Supplementary Movie**

**Legend to Supplementary tables 1-6**

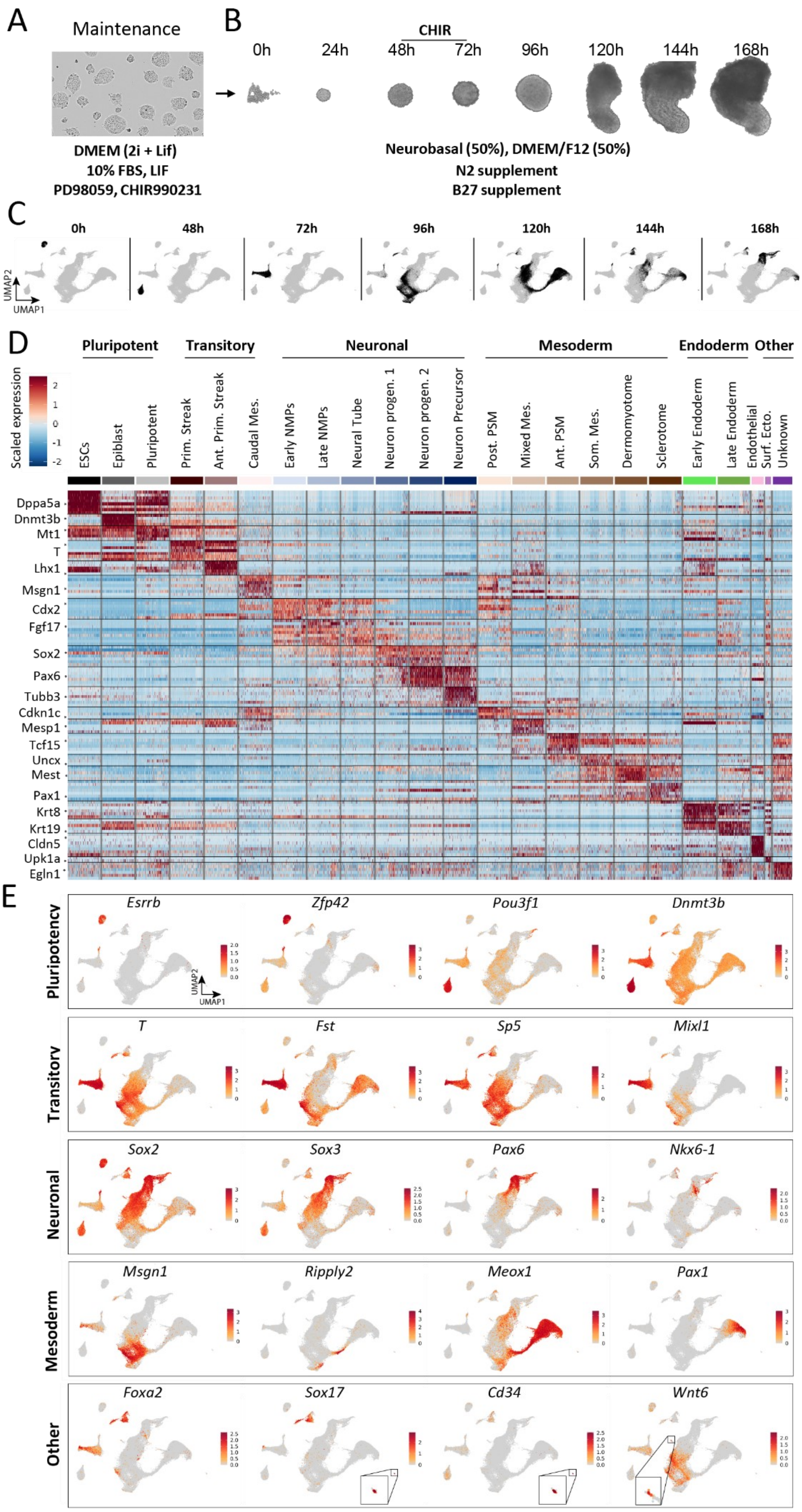


**Figure S1: Single cell RNA-seq analysis of gastruloid development (A)** Culture conditions of embryonic stem cells in this study. **(B)** Gastruloid protocol used in this study and representative image of gastruloids at each stage from aggregation to 168h. **(C)** UMAP plot highlighting, in black, cells coming from each of the time points, while the remaining cells are in grey. **(D)** Heatmap showing scaled expression of the top eight markers from each cluster identified in Fig. 1B. For visibility a random subset of 300 cells is displayed on the heatmap **(E)** UMAP plot showing the expression of the indicated marker genes.

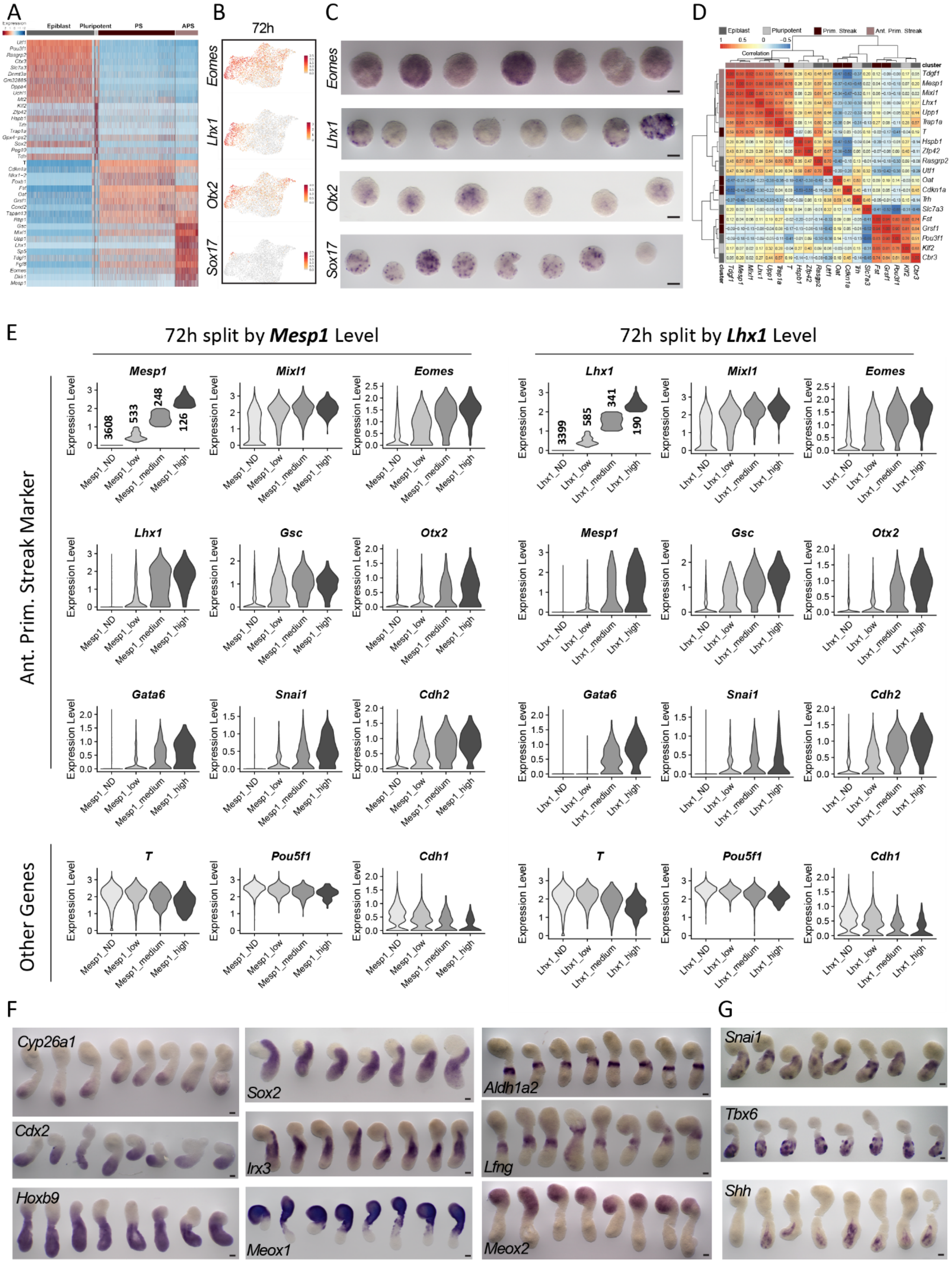


**Fig. S2: Robustness of gastruloids patterning from intrinsic variations.** **(A)** Heatmap displaying the scaled expression of the top ten markers from the clusters found at 48h and 72h. **(B)** UMAP plot of 72h gastruloid cells coloured by the normalised expression of the indicated genes. **(C)** *In situ* hybridisation of the indicated genes in 72h gastruloids. **(D)** Gene to gene Pearson correlation matrix from RNA-seq of 10 single 72h gastruloids from GSE106227. The top 5 markers of each cluster from 48h to 72h gastruloids are shown Clustering was done using WardD2 method. **(E)** Violin Plot showing the normalised expression of the indicated genes in 72h gastruloids where cells are split in four categories according to *Mesp1* (left panels) or *Lhx1* (right panels) expression level. The numbers in the first plot from each split indicate the number of cells from each category. **(F, G)** *In situ* hybridisation for indicated markers on at least seven 120h gastruloids showing robust **(F)** and less consistent **(G)** patterns of expression across gastruloids.

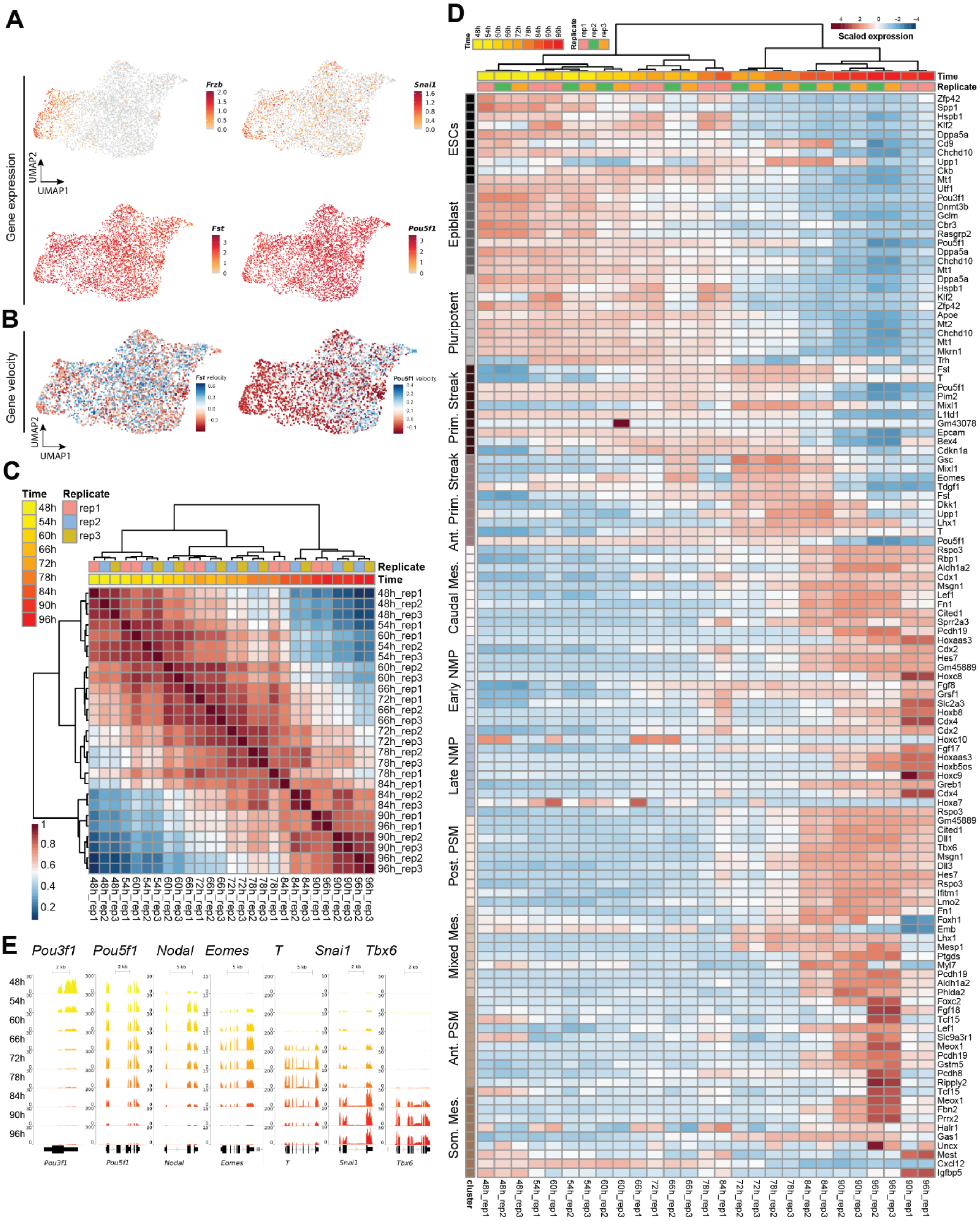

**Fig. S3: Dynamic of pluripotency exit and cell fate transitions. (A)** UMAP plot coloured according to the expression of the displayed genes in 72h gastruloids. **(B)** UMAP plot coloured by gene-specific RNA velocity on *Fst* and *Pou5f1* (coding for OCT4) in 72h gastruloids. **(C)** Spearman correlation and ward clustering of the 6h-resolution bulk RNA-seq time course on gastruloids from 48h to 96h. The 2000 most variable genes were used. **(D)** Heatmap showing the scaled log(1+ FPKM) values of bulk RNA-seq from gastruloids at 48h to 96h with a 6h resolution, and performed in three replicates. The top ten markers of the selected clusters derived from single cell RNA-seq in Fig. 1B. are shown here. **(E)** Normalised coverage of bulk RNA-seq (read per million) at the indicated loci of 48h to 96h gastruloids assessed every 6 hours, average of three replicates.

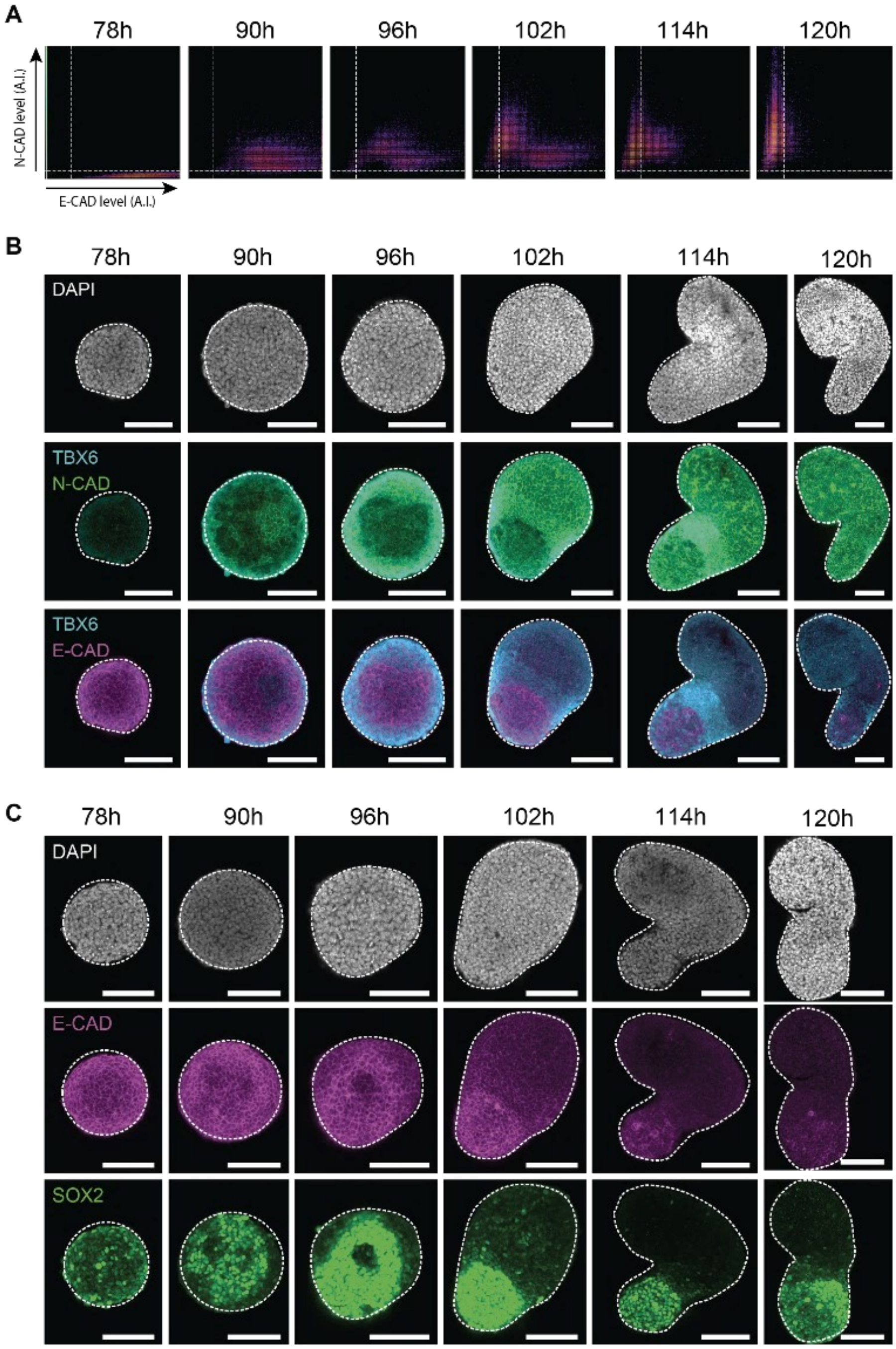


**Fig. S4: Spatio-temporal dynamic of Cadherin switch. (A)** Quantification (arbitrary unit) of membrane specific fluorescent intensity for E-Cadherin and N-Cadherin of the picture shown in **(Fig. 2E)**. **(B)** related to **Fig 2E,** Immuno-fluorescence channel for DAPI (Top), TBX6 and N-Cadherin (Middle), Tbx6 and E-Cadherin (Bottom) **(C)** related to **Fig 2F,** Confocal imaging Immuno-fluorescence showing signal for DAPI (Top), E-Cadherin (Middle) and SOX2 (Bottom). For **(A)** and **(B)**, Gastruloids contours are represented by the dashed line. z-stack is 20 µm in the gastruloids. At least 5 gastruloids were imaged per condition. Imaging was done with 20x magnification on a confocal microscope Sp8 (Leica). All compared conditions had the same illumination and acquisition settings, scale bars are 100 µm.

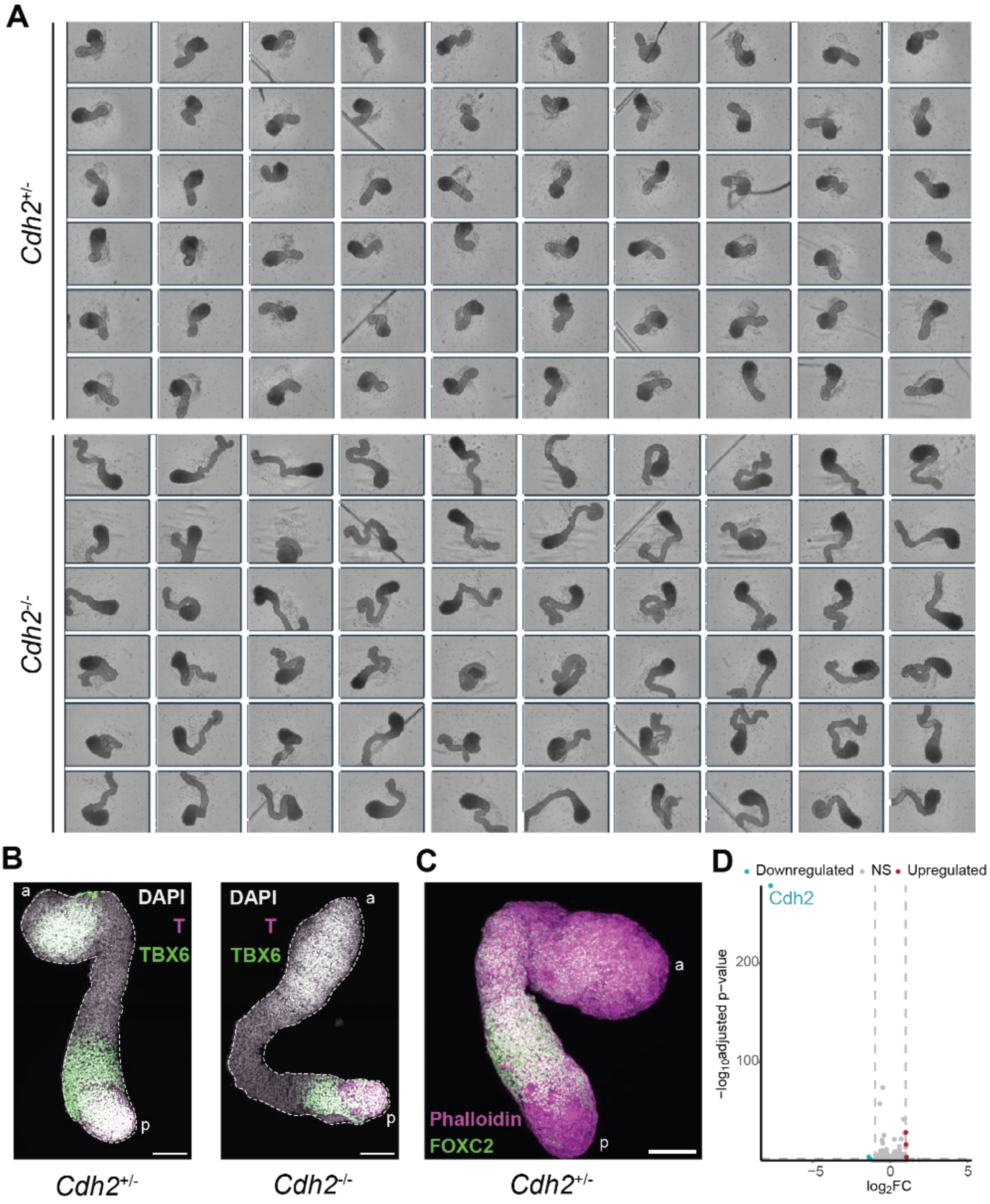


**Fig. S5: Characterisation of Cdh2 loss of function. (A)** Brightfield imaging showing example morphology of 60 gastruloids grown from *Cdh2+/-* (top panel), and *Cdh2*$^{-/-}$ (Bottom panel). **(B)** related to **Fig 3B** Maximum intensity projection of immuno-fluorescence with the DAPI signal (grey) on top of the TBX6 (green) and T/Bra (magenta) on 120h gastruloids. Scale bars are 100 µm, at least 4 gastruloids were imaged on a confocal microscope Sp8. Gastruloids contours are represented by the dashed line. **(C)** Maximum intensity projection of immuno-fluorescence for FOXC2 and F-Actin (Phalloidin) in *Cdh2*$^{+/-}$ (Het) 120h gastruloids. At least 4 gastruloids were imaged per condition. Imaging was done with 20x magnification on a confocal microscope Sp8 (Leica). All compared conditions had the same illumination and acquisition settings. scale bars are 100 µm. "a" designate the anterior side and "p" designate the posterior side. **(D)** Volcano plot showing log2 fold change over the adjusted p-value computed by DESeq2 to analyse differential gene expression between *Cdh2*$^{+/-}$ (Het) and *Cdh2*$^{-/-}$ (KO), Clone #2.

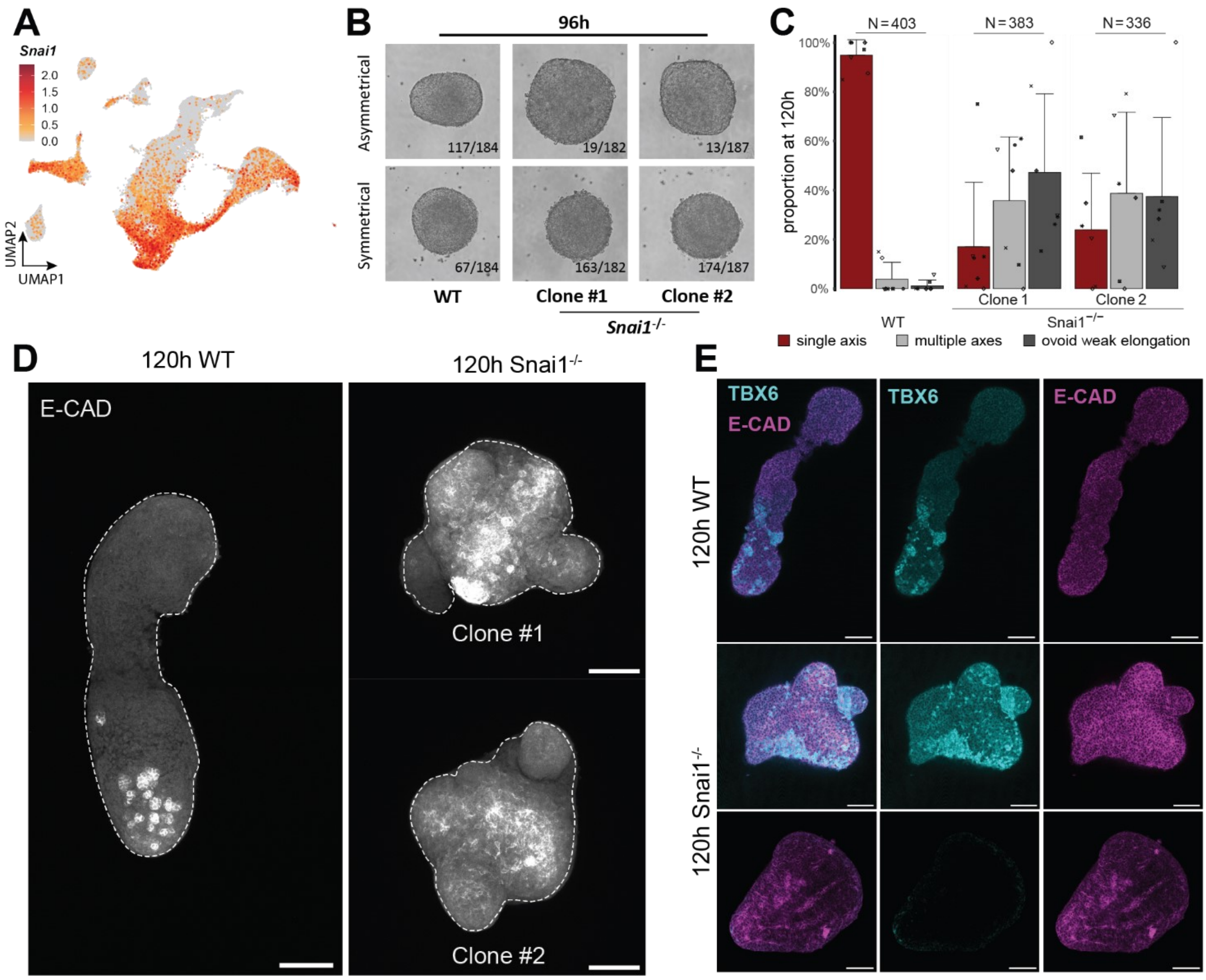


**Fig. S6: Characterization of Snai1-mediated cadherin switch function. (A)** UMAP plot coloured by *Snai1* expression in gastruloids cells from 0h to 168h (from **Fig. 1**). **(B)** Brightfield imaging showing the morphology of 96h gastruloids derived from WT and two clones of *Snai1*$^{-/-}$ ESCs. The numbers are derived from three experimental batches. **(C)** Bar chart showing proportion of morphological phenotypes observed in WT and the two clones of *Snai1*$^{-/-}$. Error bars are standard deviation to the mean. **(D)** Maximum intensity projection of immuno-fluorescence for E-Cadherin on the WT and *Snai1*$^{-/-}$ 120h gastruloids shown in **Fig. 4C**. At least 4 gastruloids were imaged on a confocal microscope Sp8 (Leica). Gastruloids contours are represented by the dashed line. **(E)** z-Section 20 µm deep of immuno-fluorescence on 120h gastruloids for TBX6 (cyan) and E-Cadherin (magenta) in wild type (top panel) and *Snai1*$^{-/-}$*gastruloids (bottom two panels)* Two different phenotypes are shown (multiple axes, middle panels) and weak elongation (Bottom panel) scale bars are 100 µm.

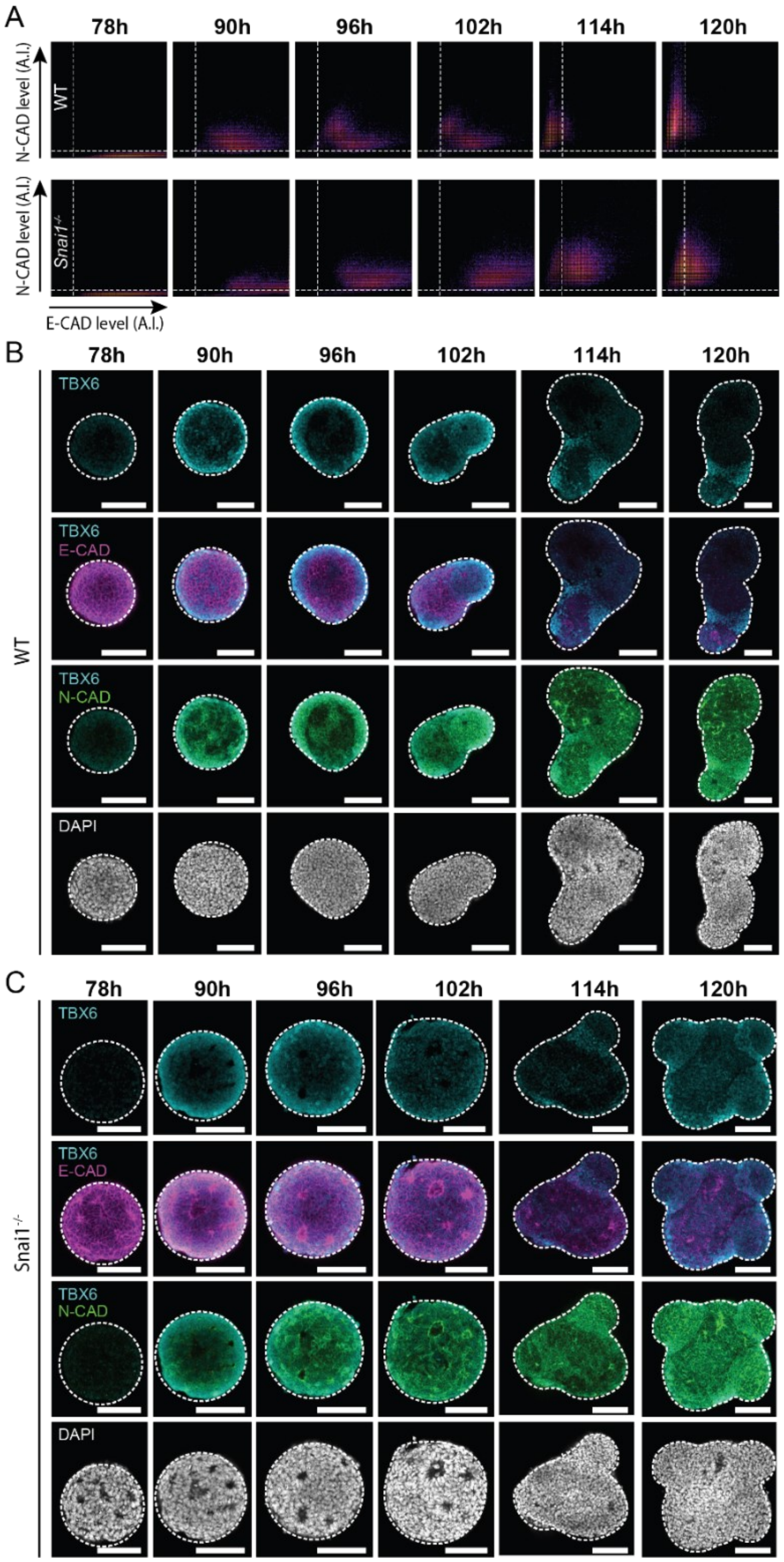


**Fig. S7: Spatio-temporal characterisation of Snai1$^{-/-}$ gastruloids. (A)** Related to **Fig. 5A** Quantification (arbitrary unit) of membrane specific fluorescent intensity for E-Cadherin and N-Cadherin in WT (top panels) and *Snai1*$^{-/-}$ (bottom panels) of the immuno-fluorescence shown in **(Fig. 5A). (B-C)** Related to **Fig. 5A,** Immuno-fluorescence signal for TBX6 (cyan) E-Cadherin (magenta), N-Cadherin (green), DAPI (grey) on WT **(B)** and *Snai1*$^{-/-}$ **(C)** gastruloids at the indicated timepoints**.** Gastruloids contours are represented by the dashed line, scale bars are 100 µm, at least 5 gastruloids were imaged on a confocal microscope Sp8 (Leica), a single stack 20 µm deep into the gastruloid is shown for each condition.

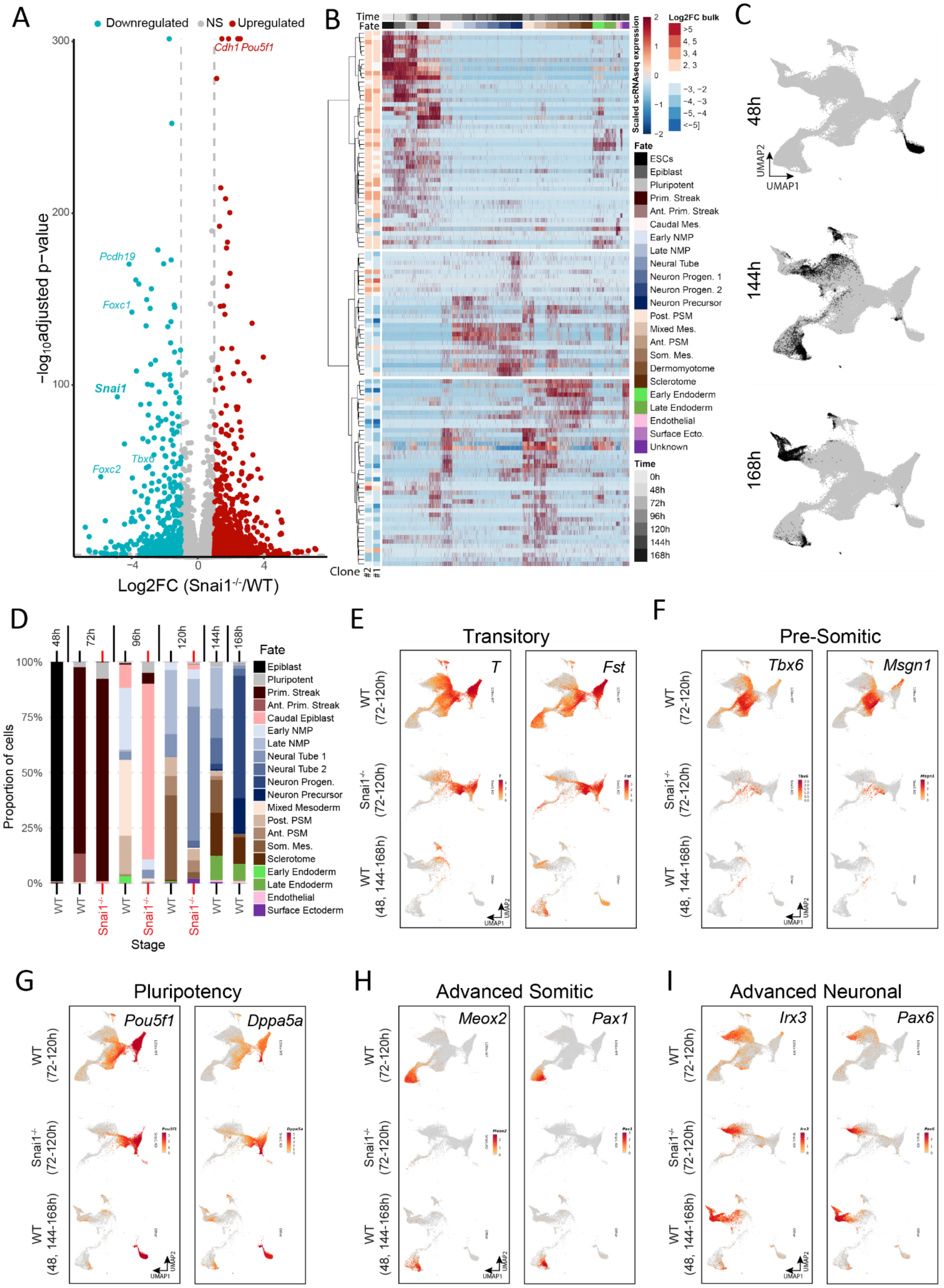


**Fig. S8: Characterization of *Snai1* mechanism of action. (See Legend on the next page)**

**Fig. S8: Characterization of *Snai1* mechanism of action. (A)** Volcano plot showing log2 fold change over the adjusted p-value computed by DESeq2 to analyse differential gene expression between Wild Type (WT) and *Snai1*$^{-/-}$ (KO) Clone #1 gastruloids at 96h. **(B)** Heatmap showing differentially expressed genes in Snai1$^{-/-}$ (abs(average log2FC) >2, p-adj<0.0005 in both clones) which are detected in single cell RNA-seq (see methods). The Clone #1 and #2 columns show genes downregulated (blue) and upregulated (red) in *Snai1*$^{-/-}$, the main heatmap corresponds to the scaled expression in a random 300cells subset from each cluster of the single cell RNA-seq time course (from Fig. 1B, C). ESCs, Embryonic Stem Cells; Prim. Streak, Primitive Streak; Ant., Anterior; Mes., Mesoderm; NMPs, Neuro-Mesodermal Progenitors; Progen., Progenitor; Surf. Ecto, Surface Ectoderm. **(C)** related to **Fig. 5G**. UMAP plot highlighting cells from WT 48h, 144h and 168h gastruloids cells (black) while the remaining cells are in grey. **(D)** Barplot coloured by cell fate proportions and split by the stage and genotype from single cell RNA-seq analysis. **(E-I)** UMAP plot coloured, by the expression of the indicated genes from WT and *Snai1*$^{-/-}$ gastruloids and the indicated times. The 48h, 144h and 168h samples were only analysed in a WT context, as such they are shown separately to allow matching samples between WT vs *Snai1*$^{-/-}$ conditions.

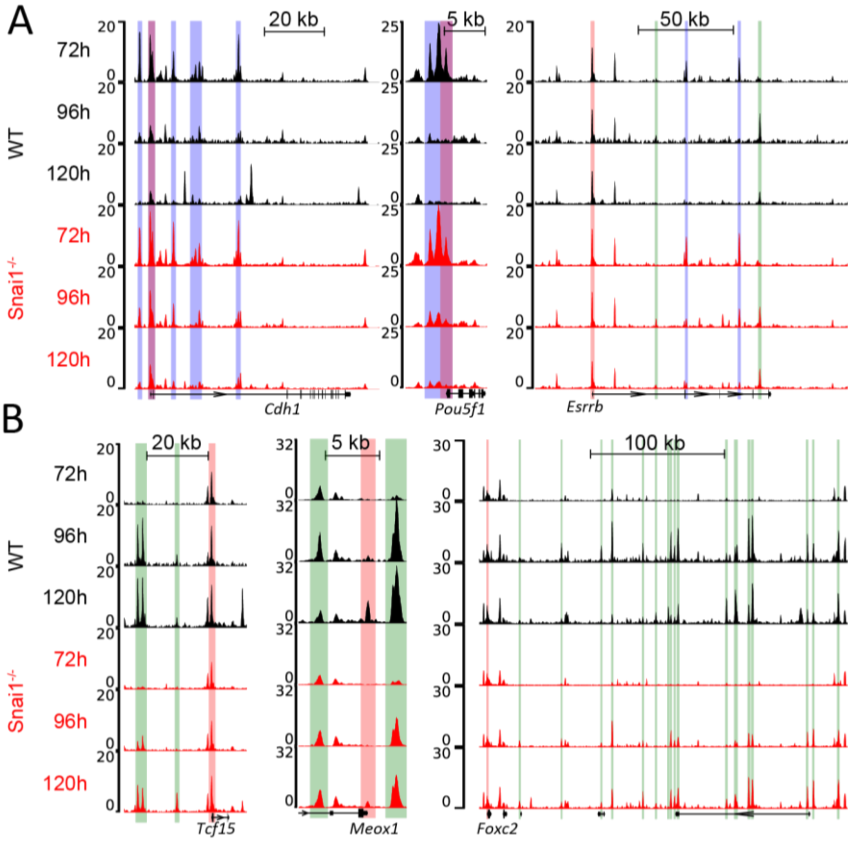


**Fig. S9: *Snai1* regulates the closing of earlier chromatin landscape. (A-B)** Normalised ATAC-seq signal (read per million read in peaks) from wild type (black) and *Snai1*$^{-/-}$ (red) gastruloids at 72h, 96h and 120h at the *Cdh1* (E-Cadherin) and the *Pou5f1* (Oct4) and *Esrrb* loci **(A)**, and at the *Tcf15*, *Meox1* and the *Foxc2* loci **(B)**. Promoter regions are highlighted in red, distal open chromatin elements (putative enhancers) which become closed **(A)** or open **(B)** at 96h are highlighted in blue and green, respectively. For *Snai1*$^{-/-}$ 72h and 96h, average of 2 clones is shown.

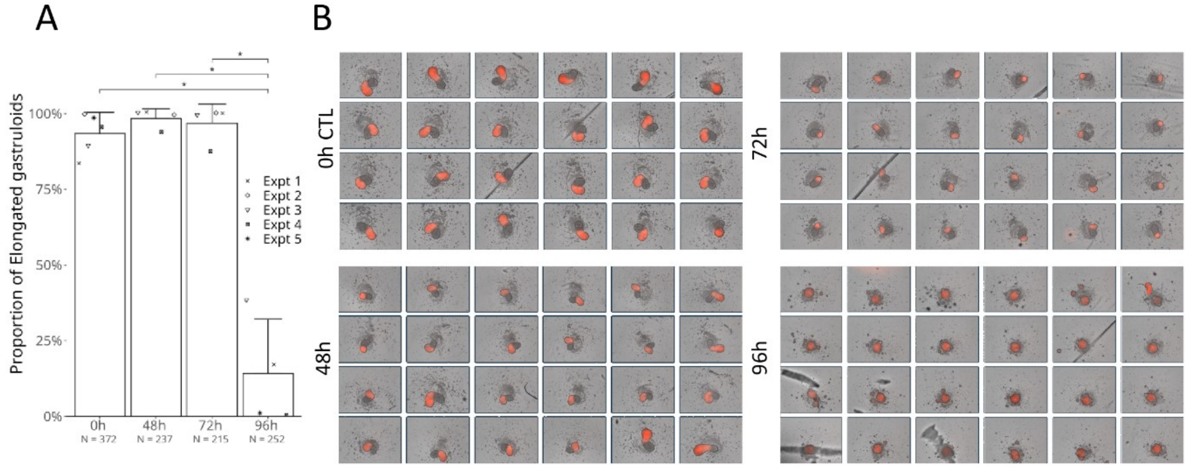


**Supplementary Note. 1: Gastruloids self-organization is resistant to single cell dissociation (A)** Proportion of elongated gastruloids when dissociated and re-aggregated at the indicated time points done on at least 4 experiments (as indicated). N represent the number of experimental batches, and n is the total number of gastruloids in each condition. *P-value* are computed from an unpaired t-test with two-tailed distribution. Error bars are standard deviation to the mean. **(B)** Representative images of 24 gastruloids for each condition. Gastruloids grown from a *Mesp2*$^{mCherry}$ reporter line were dissociated to single cells at the indicated timepoint and imaged at 120h.

**Supplementary movie 1: Gastruloid reassembly and self-organization following dissociation :** Representative movie displaying a dissociation experiment. Each frame represents 1h and gastruloids control (non-dissociated), dissociated at 48h, at 72h and at 96h are shown. The red channel derives from an H2B mCherry fluorescence driven by *Mesp2* endogenous expression

**Supplementary table 1:** List of sgRNAs used in this study.

**Supplementary table 2:** List of genotyping primers used in this study.

**Supplementary table 3:** Sanger sequencing of the ESC lines generated in this study and their associated sequencing primers.

**Supplementary table 4:** List of probes used in this study and the primers associated with their generation.

**Supplementary table 5:** List of antibodies used in this study.

**Supplementary table 6:** List of scRNA-seq samples generated in this study and whether they involved sample multiplexing using CellPlex
